# Correcting errors in PCR-derived libraries for rare allele detection by reconstructing parental and daughter strand information

**DOI:** 10.1101/2024.11.26.625438

**Authors:** Hyeonseob Lim, Soyeong Jun, Taehoon Kim, Ji Hyun Lee, Duhee Bang

**Affiliations:** Department of Chemistry, Yonsei University, 50 Yonsei-ro, Seodaemun-gu, Seoul 03722, Republic of Korea; Department of Clinical Pharmacology and Therapeutics, College of Medicine, Kyung Hee University, 26 Kyungheedae-ro, Dongdaemun-gu, Seoul 02447, Republic of Korea; Department of Biomedical Science and Technology, Kyung Hee Medical Science Research Institute, Kyung Hee University, 26 Kyungheedae-ro, Dongdaemun-gu, Seoul 02447, Republic of Korea

## Abstract

Molecular barcoding methods enable high-sensitivity detection of circulating tumor DNA that is rarely present in liquid biopsy samples. Many methods involve ligation of molecular barcodes to DNA prior to hybridization capture, enabling recovery of starting molecules. Development of polymerase chain reaction (PCR)-based methods could facilitate more cost- and labor- effective detection; however, tracking molecular identity can be difficult, as new barcodes overwrite old barcodes in each cycle. We developed a sensitive genotyping method based on a peer-to-peer network-derived identifier for error reduction in amplicon sequencing (SPIDER-seq) and enable molecular identity tracking with PCR-derived libraries using overwritten barcodes. SPIDER-seq detected mutations at frequences as low as 0.125% after only two consecutive general PCR cycles and systematically analyzed the error pattern in the peer-to-peer network. Our method could facilitate the rapid detection of mutations associated with various cancers.

## Introduction

Identifying tumor mutations can provide critical clues useful for therapy selection in cancer management ^1^. Personalized mutation profiling has been particularly effective in aiding drug selection for individual patients ^2,3^. As tumor mutations can evolve over time and thus induce relapse, early detection and subsequent continuous monitoring of minimal residual disease (MRD) is important ^4–9^. As an alternative to analysis of tissue biopsy specimens, analysis of liquid biopsy samples provides non-invasive, painless methods for identifying somatic mutations in circulating tumor DNA (ctDNA) ^10,11^. However, because ctDNA is present at very low levels in samples of cell-free DNA (cfDNA), determining whether a low proportion of observed alleles represents ctDNA or sequencing or polymerase error can be challenging ^12–14^.

Various methods have been developed recently to generate consensus sequences from molecules tagged with a unique identifier (UID) as a means of removing errors and thus improving sensitivity ^15–20^ (**Fig. 1a**). The strategy employed by these methods involves ligating a UID-containing adaptor to the starting molecules so that the resulting amplified daughter molecules can be grouped according to the UID. A consensus sequence is generated from the grouped molecules, and error molecules, which typically constitute a low percentage of the total, can then be removed. Combined with hybridization capture ^21^, these methods have been employed to profile somatic mutations in dozens of genes. However, hybridization capture is time consuming (requiring 2–3 days) and costly ^22,23^. In addition, the method provides an on-target ratio of only 20–30% when targeting ∼200 genes^24^. This strategy is thus inefficient for repetitive inspection of only a few hotspots, as is the case with MRD monitoring. A promising alternative approach is the identification of mutant DNA using droplet digital PCR; however, considerable optimization is required for probe preparation, and the method is not ideal for multiplex detection ^25–28^. More rapid and cost-effective methods are needed for personalized monitoring of multiple target mutations ^2,29^.

**Fig. 1.**
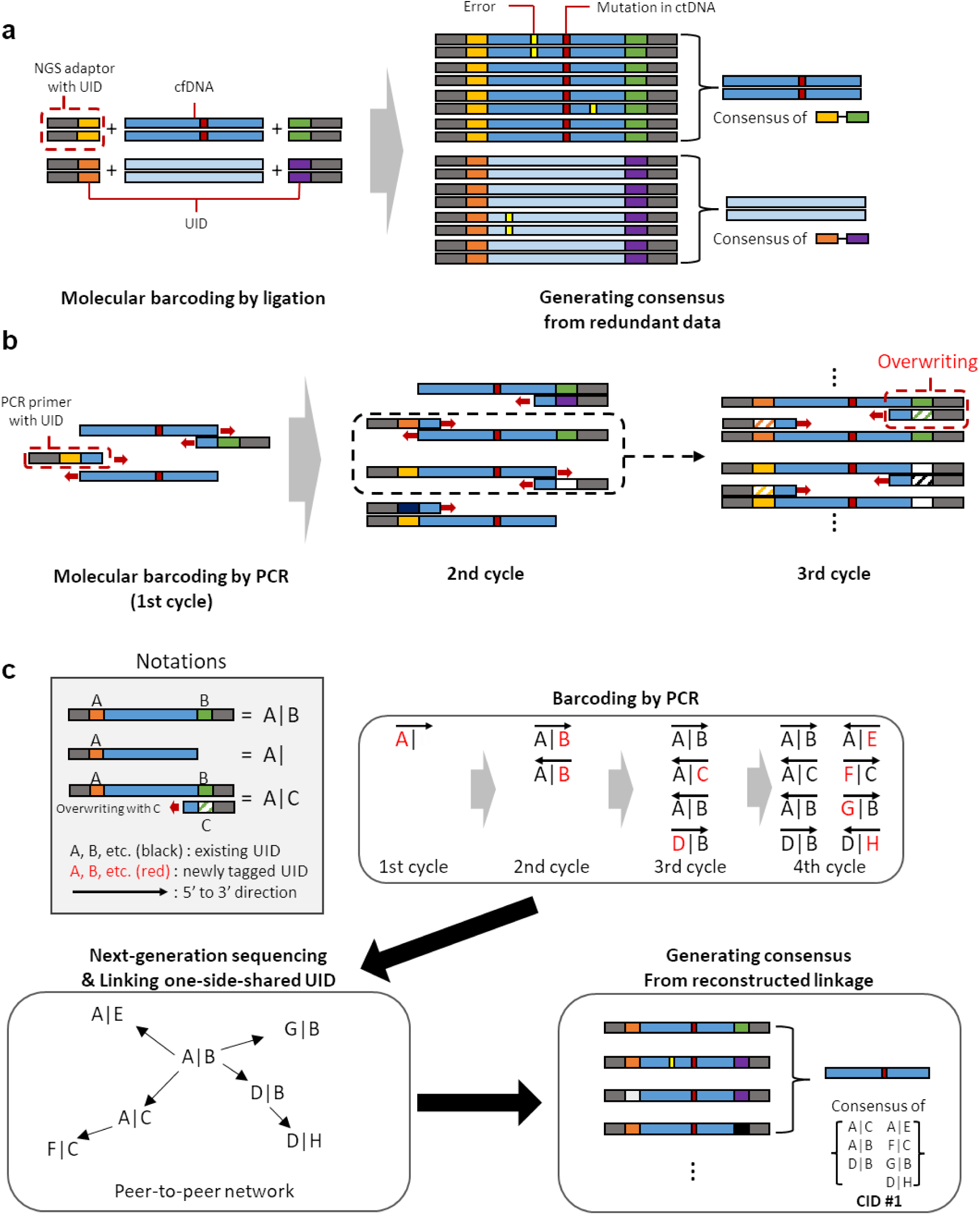
Schematic illustration of molecular barcoding systems. (**a**) Example of a simplified ligation-based barcoding system. UIDs attached by ligation ensure the original molecules’ identity in this system. (**b**) When incorporating UIDs via PCR primers, UIDs are overwritten during repeated PCR cycles. (**c**) Creating a linkage between the two strands using the shared UID. The small red block in the sequence indicates a nucleotide variant, and the small yellow block in the sequence indicates a polymerase or sequencing error introduced during the preparation steps. UID, unique identifier; CID, cluster identifier.

We hypothesized that amplicon sequencing would be more efficient for personalized monitoring of multiple target mutations because the assays can be completed rapidly and easily with a high on-target ratio using inexpensive PCR reagents. Amplicon sequencing is also suitable for low amounts of input material. However, molecular tagging with a UID in PCR is more complex than ligation reactions. When incorporating a UID sequence via PCR primers, the UID sequences are overwritten over repeated PCR cycles (**Fig. 1b**); thus, multiple UID-pairs must be generated from the starting molecule. To the best of our knowledge, only two types of methods currently enable molecular tagging of an amplicon library. One type of method involves limiting the number of amplifications (e.g., 2–3 PCR cycles) to prevent overwriting of the UID ^30–32^. However, restricting the number of PCR cycles increases the difficulty of preparing libraries of inhibitor-treated clinical samples. The second type of method involves the use of linear amplification followed by single-stranded ligation for attaching the UID ^23^. Although this type of method enables detection of mutant alleles with a frequency of <0.01%, the complex experimental process, which includes a 24-hour ligation and three rounds of PCR, makes the approach laborious.

To overcome the abovementioned drawbacks, we evaluated the suitability of a general PCR approach. The recent development of blockchain technology ^33^ motivated us to combine multiple UID-pairs generated from each individual starting molecule strand into a single identity. Blockchain technology employs a distributed networking strategy to record information so that the resulting data can be identified by integrating all individual records searched through a peer-to-peer network. Similarly, integrated information can be obtained from peer-to-peer networks constructed using all daughter molecules derived from the first-copied strand from the original molecule. The daughter strand resulting from each amplification cycle with a UID-containing primer will contain two UIDs. The first is the overwritten UID incorporated by the primer, and the other is the replicated UID derived from the parental strand. Both the parental and daughter strands thus share one UID (**Fig. 1b**). We hypothesized that we could create a link between the two strands using the shared UID and that the created linkage could be extended to the granddaughter strand. The overwritten UID in the daughter strand could be used again as a shared UID with a granddaughter strand so that a connection could be achieved with all descendant strands derived from the first-copied strand (**Fig. 1c**). The UIDs in the network could therefore be considered a cluster (i.e., grouped UIDs) that can serve as an integrated identifier (designated a cluster identifier [CID]). We also hypothesized that the rate of sequencing and polymerase errors could be reduced if the consensus sequence was generated based on the CID.

In addition to building a consensus sequence, we expected that the linkage could be used to construct a lineage of amplification. By placing the first-attached UID from a cluster in the top position, daughter strands can be appended in order of generation similar to a rooted phylogenetic tree. This arrangement can be used to characterize the error pattern by investigating the continuity of errors along the branch. Sporadic errors, such as sequencing errors, should be focused at nodes, whereas polymerase errors should be conserved along the branch. We postulated that this pattern could be used to review the effectiveness of error correction efforts.

In the present study, we developed a sensitive genotyping method based on a peer-to-peer network-derived identifier for error reduction in amplicon sequencing (SPIDER-seq). Errors are reduced by generating consensus using CID based on the amplicon sequencing method. Using SPIDER-seq, we demonstrated proof-of-concept using a model oligonucleotide and a set of mock ctDNA references. We clustered daughter strands by constructing a peer-to-peer network and demonstrated that generating CID-based consensus effectively reduces errors, and the sensitivity of the approach enabled the detection of a 0.125% ctDNA allele frequency (AF) with high accuracy and reproducibility. By determining the lineage of amplification, we found that the majority of sequencing errors were corrected, but polymerase errors introduced in the early cycles of amplification were not corrected. We also demonstrated that SPIDER-seq can be applied to multiple targets via multiplex PCR ^34,35^.

## Materials and Methods

### Materials

We designed and obtained a custom-produced oligonucleotide (Ultramer™ oligo with standard desalting purification from Integrated DNA Technologies) for use in model experiments to demonstrate SPIDER-seq performance. The oligonucleotide was designed to mimic the genomic sequence containing the BRAF p.V600E mutation and was 173 nt in length to simulate the typical size of plasma-derived cfDNA. To distinguish DNA molecules, genomic sequences were replaced with a 12-nt sequence of degenerate random bases (hereafter referred to as barcode content) (**Supplementary Table 1**). For experiments designed to demonstrate the feasibility of SPIDER-seq for detecting ctDNA, we utilized Seraseq^TM^ ctDNA Mutation Mix v2 (Seracare), a mock cfDNA reference spanning a variant allele frequency of 0–1% (**Supplementary Table 2**). Details regarding the allele frequency of each locus and concentration are provided in the manufacturer’s information.

### Design of PCR primers

Given the short average length of cfDNA, we designed primers to target an approximately 100-bp region of the target gene to facilitate amplification. At the 5’ end of the designed primer, a sequencing adaptor, flanking sequence, and UID comprised of 16 degenerate and 4 fixed bases (NNNNXNNNNXNNNNXNNNNX, where N=degenerate base and X=fixed base) were concatenated. The flanking sequence and fixed bases in the UID sequences were designed to have different sequence combinations to ensure sequencing quality control. The sequences of all designed primers are listed in **Supplementary Table 1**. All primers were synthesized by Integrated DNA Technologies (oligos with standard desalting purification).

### UID incorporation and preparation of library for sequencing

The library was constructed via two rounds of PCR amplification. The first round of amplification was conducted to incorporate the UID sequence. For model experiments, 100 μM oligo was diluted 10^6^-fold to restrict the number of molecules and then used as a template. Recipes and cycling conditions for the first-round PCRs were as follows.

PCR recipe using KAPA HiFi polymerase: starting material, 1 μl of forward primer (10 μM), 1 μl of reverse primer (10 μM), 4 μl of 5× KAPA HiFi buffer, 0.6 μl of dNTPs (10 mM each), 0.4 μl of KAPA HiFi HotStart polymerase, and nuclease-free water to a final volume of 20 μl.

PCR recipe using the QIAGEN Multiplex PCR kit: starting material, 1 μl of forward primer (10 μM), 1 μl of reverse primer (10 μM), 10 μl of 2× QIAGEN Multiplex PCR Master Mix, and nuclease-free water to a final volume of 20 μl.

PCR recipe using Phusion High-Fidelity DNA polymerase: starting material, 1 μl of forward primer (10 μM), 1 μl of reverse primer (10 μM), 4 μl of 5× Phusion HF buffer, 0.4 μl of dNTPs (10 mM each), 0.2 μl of Phusion DNA polymerase, and nuclease-free water to a final volume of 20 μl.

PCR conditions using KAPA HiFi polymerase: 95°C for 3 min, followed by 6 cycles of 98°C for 20 s, 56°C for 15 s, 72°C for 30 s; and 72°C for 1 min.

PCR conditions using the QIAGEN Multiplex PCR kit: 95°C for 15 min, followed by 6 cycles of 94°C for 30 s, 56°C for 90 s, 72°C for 1 min; and 72°C for 10 min.

PCR conditions using Phusion High-Fidelity DNA polymerase: 98°C for 30 s, followed by 6 cycles of 98°C for 10 s, 56°C for 15 s, 72°C for 30 s; and 72°C for 5 min.

For experiments using mock cfDNA reference and targeting a single gene (*BRAF*), 1 μl of reference DNA was used as the starting template, corresponding to 3,697–4,788 haploid genome equivalents (hGEs) (**Supplementary Table 3**).

PCR recipe using KAPA HiFi polymerase: starting material, 1 μl of forward primer (10 μM), 1 μl of reverse primer (10 μM), 4 μl of 5× KAPA HiFi buffer, 0.6 μl of dNTPs (10 mM each), 0.4 μl of KAPA HiFi HotStart polymerase, and nuclease-free water to a final volume of 20 μl.

PCR conditions using KAPA HiFi polymerase: 95°C for 3 min, followed by 8 cycles of 98°C for 20 s, 56°C for 15 s, 72°C for 30 s; and 72°C for 1 min.

For experiments using mock cfDNA reference and targeting multiple genes, 2 μl of reference DNA was used as the starting template, corresponding to 8,424–9,576 hGEs (**Supplementary Table 3**).

PCR recipe using the QIAGEN Multiplex PCR kit: starting material, 1 μl of forward primer mixture (10 μM), 1 μl of reverse primer mixture (10 μM), 10 μl of 2× QIAGEN Multiplex PCR Master Mix, and nuclease-free water to a final volume of 20 μl.

PCR conditions using the QIAGEN Multiplex PCR kit: 95°C for 15 min, followed by 8 cycles of 94°C for 30 s, 56°C for 90 s, 72°C for 1 min; and 72°C for 10 min.

After the first-round amplification, the product was used directly used in the next step without purification in order to prevent loss of product molecules. A total of 8 separate 50-μl reactions were carried out using 2.5 μl of product from the first-round amplification. The PCR recipe was as follows: 2.5 μl of product from the first-round amplification, 2.5 μl of NEBNext i5 primer (10 μM), 2.5 μl of NEBNext i7 primer (10 μM) (NEB), 5 μl of 5× KAPA HiFi buffer, 0.75 μl of dNTPs (10 mM each), 0.5 μl of KAPA HiFi HotStart polymerase, and nuclease-free water to a final volume of 50 μl. Amplification was performed under the following conditions: 98°C for 30 s, followed by 20–22 cycles of 98°C for 10 s, 65°C for 30 s, 72°C for 30 s; and 72°C for 5 min. The combined product (∼ 300 bp) was size selected on agarose gel and then purified using a MinElute Gel Extraction Kit (Qiagen). The products were sequenced on an Illumina NovaSeq 6000 or NextSeq 500 platform. The number of paired reads in the sequencing data for each sample is provided in **Supplementary Table 4.**

### Trimming of raw data

Primer sequences were trimmed from the raw data, and the UID sequence of each read was identified from the primer region. To minimize misidentifications of UID sequences, low-quality reads meeting the following conditions were filtered out: (i) average phred quality <30; (ii) low-quality UID sequence in which the fixed bases differed from the designed sequence or the minimum phred quality of the UID bases was <25; (iii) high-GC UIDs in which the G-C proportion was ≥0.8. During analysis of the barcode contents of synthesized oligonucleotides, reads with an incorrect flanking sequence near the barcode content were also filtered out. In analyses of data from experiments using cfDNA reference, the trimmed data were aligned to the reference genome (hg38) using BWA-MEM (Ver: 0.7.15) ^36^. The aligned data were converted to BAM format and indexed using SAMTOOLS (ver. 1.9) ^37^. Reads exhibiting a mapping quality <55 or that were mapped with soft-clipping were also filtered out. The remaining reads were subjected to subsequent steps. If necessary, some data were down-sampled using seqtk (available at https://github.com/lh3/seqtk) in the raw data state and then used for downstream analyses.

### Clustering via construction of a peer-to-peer network

To construct a peer-to-peer network, the UID-pairs for each molecule were first organized. Paired-UIDs with the same left or right UID were grouped to create linkages between UID-pairs. Improper UIDs in which the number of paired-UIDs was equal to or greater than the number of PCR cycles were removed. Beginning with addition of a randomly selected UID to the list of clusters, paired-UIDs of existing UIDs were added to extend the elements. Paired-UID addition was conducted recursively until no more paired-UIDs to be added remained. The cluster was then checked to confirm the absence of more than the possible number of UIDs (i.e., 2^cycle-2^, **Supplementary Note**) and determine if multiple paths existed between any two UIDs (designated ‘multibridge’). If either case occurred, the cluster was not used to create consensus. The list of UIDs was then designated a CID, and the read IDs supporting the CID were saved into a mapping file, which was used to designate the CID of each read in the BAM-formatted data.

### Analysis of barcode contents

After constructing the peer-to-peer network, barcode contents were analyzed using the trimmed fastq data. The barcode content of each read was identified based on regular expression and then collected according to the CID. Clusters in which there was only one supporting paired read were excluded. If one or two sequence mismatches between the major barcode and other barcodes was observed among barcode contents in the same cluster, the barcode contents were corrected so as to be the same as the major barcode. The specificity of the major barcode was then calculated.

### Lineage construction using cluster information

The major UID in a particular cluster (the UID with the largest number of paired-UIDs) was considered the first-tagged UID (i.e., the origin UID). Subsequent linked UIDs were then ordered next to the existing UIDs using a depth-first search algorithm. After all paths were completed, a phylogenetic tree was generated using UID as the vertex and the relationship between linked UIDs as the edge. Tree data were visualized as a dendrogram using the networkD3 package (available at https://CRAN.R-project.org/package=networkD3). To facilitate ease of computing, a peer-to-peer network with a UID-to-UID structure was constructed. In the visualization process, the structure reverted to a strand-to-strand–based tree.

### Analysis of cfDNA reference standards

To analyze substitution mutations, reads from the aligned data were parsed using the pysam module of python, and target bases were identified using the get_reference_sequence function of pysam. The consensus base of each targeted position was then determined per each CID. Clusters for which there were <2 paired-reads (i.e., total of 4 reads), the size (i.e. number of left UIDs + number of right UIDs) was <3, or the frequency of major bases was <0.7 were excluded. The number of consensus bases supporting A, T, C, and G was then determined.

For indel analyses, target mutations were listed in vcf format, which could be obtained using indel caller (e.g., VarDict) or by manual scripting. To determine whether indel mutations were present in the reads, query strings corresponding to the mutated and wild-type sequences were searched within the read sequence. To create query sequences, sequences consisting of 10 bp upstream and downstream were attached to the wild-type or mutant sequence. The genotype of each read was then categorized as either indel or wild-type, and the major genotype per CID was determined and designated the consensus genotype. Clusters for which there were <2 paired-reads (i.e., total of 4 reads), the size (i.e., number of left UIDs + number of right UIDs) was <3, or the frequency of the major genotype was <0.7 were excluded.

### UID incorporation and preparation of the library for hybridization capture

A 2 μl sample of cfDNA reference standard (7,394–9,576 hGEs, **Supplementary Table 3**) was end-repaired and A-tailed using 5× ER/A-tailing Enzyme Mix (Enzymatics). The NEBNext Adaptor for Illumina (NEB) was then ligated to the DNA ends using WGS ligase (Enzymatics), and the resulting product was digested using USER enzyme (NEB).

The products were indexed with custom-designed i5 and i7 primers (**Supplementary Table 1**). Of the eight index bases, five were used for the UID, and the remaining three bases were used for the sample barcode. Four indices each were designed for i5 and i7 and synthesized by Integrated DNA Technologies. Indexing was performed by PCR under the following conditions. The ligated product was mixed with 2.5 μl of custom i5 primer (10 μM), 2.5 μl of custom i7 primer (10 μM), 5 μl of 5× KAPA HiFi buffer, 0.75 μl of dNTPs (10 mM each), 0.5 μl of KAPA HiFi HotStart polymerase, and nuclease-free water to a final volume of 50 μl. PCR cycling was programmed as follows: 98°C for 30 s, followed by 8 cycles of 98°C for 10 s, 65°C for 30 s, 72°C for 30 s; and 72°C for 5 min. The product was purified using 1.2× Ampure XP beads (Beckman Coulter). Finally, hybridization capture was performed by Celemics (Korea), and the product was sequenced on an Illumina NovaSeq 6000 platform.

### Analysis of hybridization capture samples

The data were first demultiplexed using 3-bp sample barcodes in the i5 and i7 indexes, after which the UID sequence was extracted from the indexes. Similar to the quality trimming step in the amplicon sequencing analysis, low-quality reads meeting the following conditions were filtered out: (i) average phred quality <30; (ii) high-GC UIDs for which the G-C proportion was ≥0.8. The filtered data were mapped to hg38 using BWA-MEM; reads with a mapping quality <55 or those mapped with soft-clipping were also filtered out.

Information regarding paired-UIDs was collected per each genomic coordinate having the same start and end position, and the cluster was then constructed using these genomic coordinates. Excepting the construction of clusters using only reads with identical start and end positions, the clustering and consensus base creation processes were the same as that used for analysis of the amplicon library.

### Statistics and reproducibility

To compare differences between the groups, the Wilcoxon rank sum test was used in Figure 2e and the Wilcoxon signed rank test was used in Figure 3b and d and Supplementary Figures 11c, 14a, 15b.

**Fig. 2.**
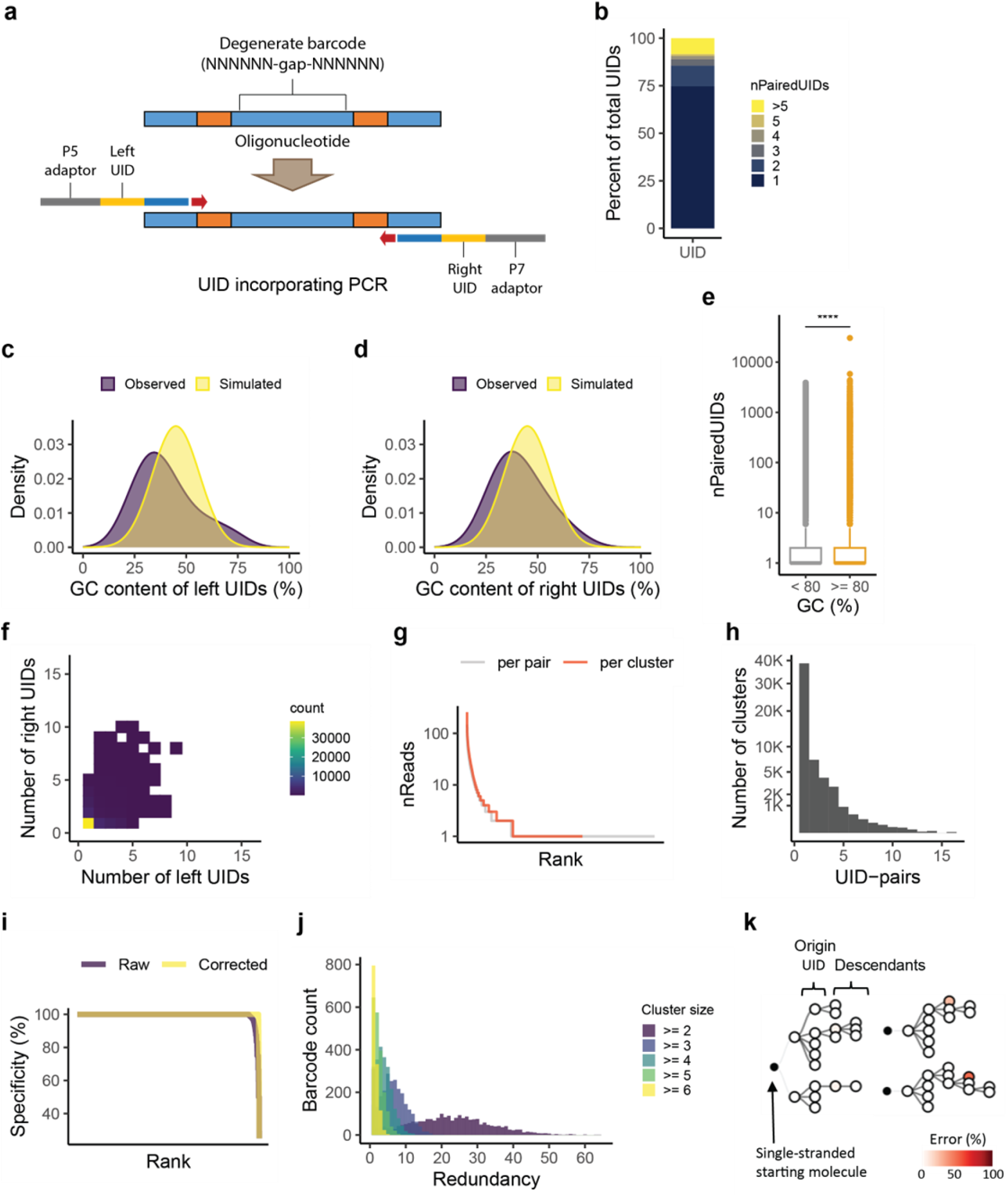
Model experiment demonstrating feasibility of cluster construction. (**a**) Schematic illustration of the experiment. The oligonucleotide was designed to contain a 12-nt barcode for molecular identity. Primers were designed to have a UID and adaptor sequences for Illumina sequencing. (**b**) Number of paired-UIDs (nPairedUIDs). (**c, d**) GC content (%) of left UIDs (**c**) and right UIDs (**d**). Simulated data was created using a pool of 100,000 randomly generated UID sequences. (**e**) Comparison of nPairedUIDs between normal-GC (<80%) and high-GC (≥80%) groups. Groups were compared using the two-sided Wilcoxon rank sum test. (****, *P* = 2.50 × 10^−152^). (**f**) Cluster size distribution. (**g**) Number of reads per UID-pair and per cluster, in which pairs and clusters are shown in ranked order. (**h**) Distribution of UID-pairs per cluster. (**i**) Specificity (%) of clusters before and after barcode content correction within a hamming distance of 2, in which clusters are shown in ranked order. (**j**) Distribution of redundancy for a given cluster size. (**k**) Representative lineages of clusters in which sequencing errors were observed.

**Fig. 3.**
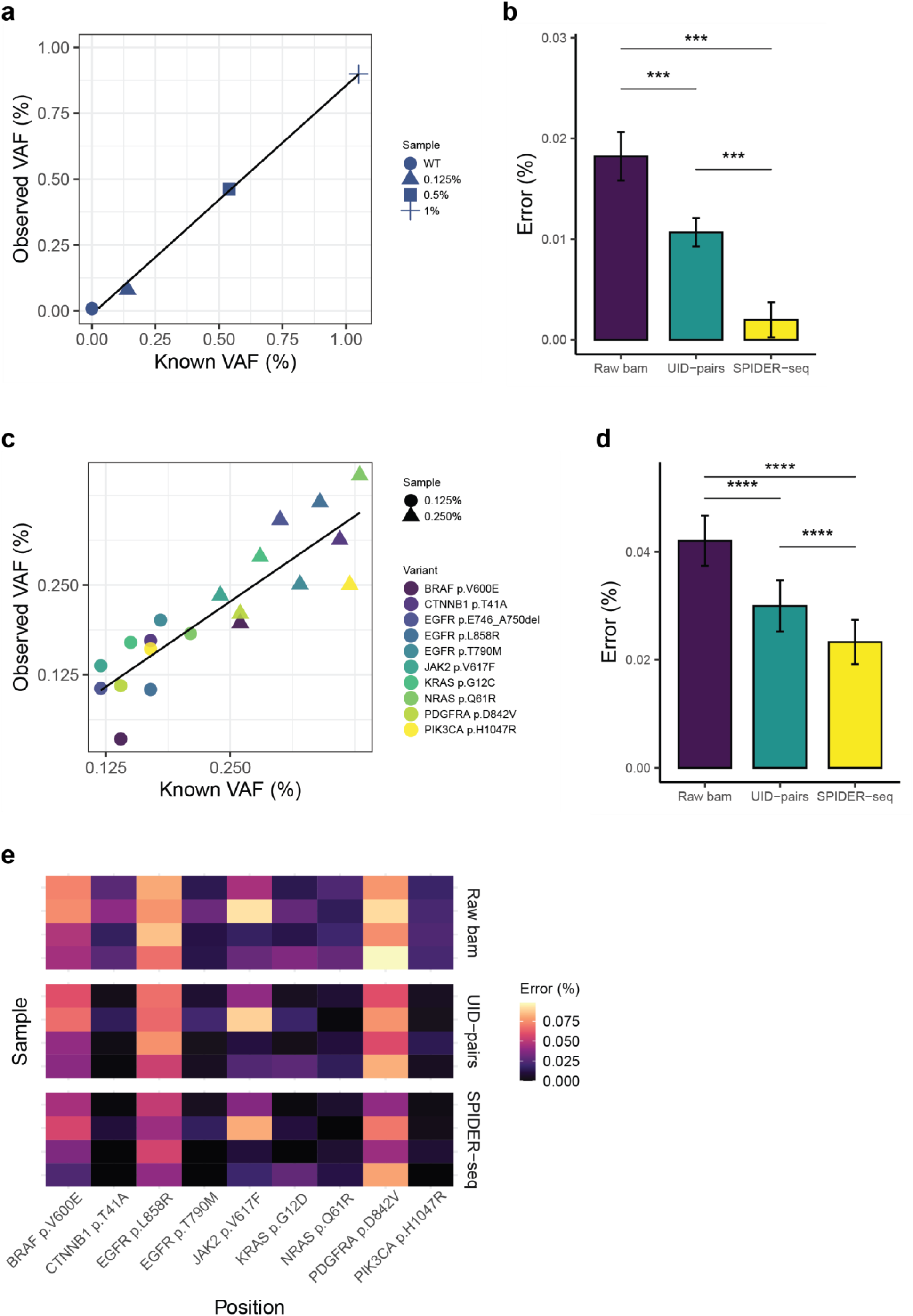
Performance of SPIDER-seq in detecting a single mutation (A and B) and multiple mutations (C–E). (a) Comparison of VAFs observed using SPIDER-seq with known VAFs provided by the manufacturer. For each sample, the mean observed VAF from replicate experiments is presented. The line is a linear fit. Pearson *r* = 0.99871. (**b**) Comparison of error rates (%) across the following methods: base count from raw bam files, base count using UID-pairs, and base count using clusters (SPIDER-seq). Error bars represent standard error of the mean. Methods were compared using the Wilcoxon signed rank test (*P* = 4.88 × 10^−4^ for all two-group comparisons). (**c**) Comparison of VAFs observed using SPIDER-seq with known VAFs provided by the manufacturer. For each sample and variant, the mean observed VAF from replicate experiments is presented. The line is a linear fit. Pearson *r* = 0.881145. (**d**) Comparison of error rate (%) across the following methods: base count from raw bam files, base count using UID-pairs, and base count using clusters (SPIDER-seq). Error bars represent the standard error of the mean. Methods were compared using the Wilcoxon signed rank test (*P* = 2.91× 10^−11^ between raw bam and UID-pairs, *P* = 2.91× 10^−11^ between raw bam and SPIDER-seq, *P* = 2.91× 10^−10^ between SPIDER-seq & UID-pairs). (**E**) Error rate (%) across positions. Error rates were calculated by dividing the number of non-reference bases by the total number of bases at the targeted position. The indel position was excluded from the error rate calculation because non-reference bases cannot be counted.

## Results

### Feasibility of constructing a cluster based on a peer-to-peer network

To assess the feasibility of constructing a cluster based on a peer-to-peer network, we conducted a model experiment using an oligonucleotide containing a barcode consisting of 12-nt degenerate bases. The oligonucleotide was then amplified along with a pair of UID-containing primers via six rounds of thermal cycling using KAPA HiFi polymerase (**Methods**, **Fig. 2a**). An amplicon library was then prepared, and paired-end sequencing was performed. We hypothesized that the barcode content would be identical across the reads in a single CID obtained by linking the UID-pairs.

We initially investigated the characteristics of the UID-pairs using the sequencing data. Assuming an ideal case in which each strand is used repeatedly as a template across cycles, each strand would be expected to produce multiple daughter strands in the amplification experiment, such that the parental strand could be linked directly to the multiple daughter strands (**Supplementary Fig. 1**). The possible number of daughter strands obtained from each parental strand was estimated at a maximum of five, assuming that the first-copied strand was synthesized in the first amplification cycle and daughter strands would be produced in the second through sixth cycles. Any parental strand synthesized later than the first cycle would therefore produce fewer than five daughter strands. In other words, a UID could have a maximum of five paired-UIDs from different daughter strands. As expected, we found that most UIDs had five or less paired-UIDs.

Although most UIDs had no more than five paired-UIDs, 8.41% of UIDs produced more than five paired-UIDs (**Fig. 2b**). We concluded that those UIDs linked to more than five UIDs were associated with high melting temperatures of the high-GC sequences. Graphs of the distribution of observed GC content exhibited a distinct right tailing indicative of high GC content (**Fig. 2c, d**), which was not observed in an ideal distribution of a randomly generated UID set. We presume that a primer with a high-GC UID may preferentially reattach, leading to the initiation of a new lineage independent of the original one. We also found that more daughter-UIDs tended to be produced from parental UIDs with a GC content of ≥80% (**Fig. 2e**). As cases involving more than five paired-UIDs could result in over-collapsing, false consensus would be obtained. In particular, the sensitivity of detecting mutations in ctDNA would decrease if UID-pairs derived from normal DNA were collapsed to UID-pairs derived from ctDNA. Therefore, we filtered out UIDs for which the number of paired-UIDs was higher than the number of cycles or cases in which the GC content was ≥80%.

Parent and daughter strands were then linked via the peer-to-peer network (**Supplementary Fig. 2**). Extension of the linkage between strands was carried out in a manner similar to *de novo* assembly ^38,39^. In order to simplify the computational process, individual UIDs were used as vertexes (**Clustering via construction of a peer-to-peer network in the Materials and Methods**). Starting from a seed UID randomly selected from the observed UIDs, paired-UIDs were recursively added to the opposite site of the strand until no new paired-UIDs to add remained. The list of paired-UIDs was considered a cluster, and a CID was assigned to each cluster. This process resulted in the formation of 58,114 clusters consisting of various sets of UID-pairs (**Fig. 2f**). For each cluster, the UIDs on each side (left and right sides of the amplicon, designated “left UID” and “right UID”) were used in a balanced manner; the maximum total number of left and right UIDs per cluster (i.e., number of left UIDs + number of right UIDs, designated “cluster size”) was 37.

The clusters were characterized by first determining how many reads and UID-pairs supported each CID. On average, each CID was composed of 6.283 paired-reads (**Fig. 2G**). In contrast, fewer reads (2.955 on average) supported each UID-pair than each cluster. This suggests that using CIDs is more advantageous than using UID pairs, by leveraging more reads to generate consensus on error reduction. In terms of UID-pairs, clusters with a size of 2 that consisted of only one UID-pair accounted for 66.05% of all clusters (**Fig. 2h**). The remainder of all clusters (33.95% of the total) were supported by 95,920 UID-pairs (68.94% of all UID-pairs). The majority of UID-pairs were therefore linked with the other UID-pairs.

To assess the accuracy of cluster construction, the specificity of the barcode content per cluster was determined by calculating the frequency of major barcode content per cluster. As expected, most of the clusters contained the same barcode content regardless of cluster size (**Fig. 2i**). Even in cases in which the specificity was not 100%, the barcode content sequences were very similar, and only 1 to 2 content mismatches were observed. After correction of these mismatches, 99.09% of the clusters exhibited identical barcodes (i.e., specificity of 100%). These mismatches were assumed to have arisen during PCR and sequencing.

We then determined how many clusters are generated from a starting molecule. In PCR assays, one starting molecule can produce a copy molecule (i.e., first-copied strand) in each cycle. Thus, multiple clusters can be initiated from a single molecule in every PCR cycle (**Supplementary Fig. 1**). Theoretically therefore, a maximum of five clusters could be generated from a template oligonucleotide through six amplification cycles. As expected, most of the barcode contents were redundantly observed in multiple clusters (**Fig. 2j**). However, some barcode contents were observed in more than five clusters. This was attributed to breakage of the clusters into multiple pieces due to missing UID-pairs during the purification or sequencing steps (**Supplementary Fig. 3**). This cluster breakage was also considered to be the cause of the high proportion of clusters with a size of 2 (**Fig. 2h**). The redundancy actually decreased if we discarded the clusters with a size less than 3. Importantly, we hypothesize that the redundant clusters would be helpful in attempts to detect extremely low amounts of ctDNA because multiple consensus can be obtained from one molecule.

### Use of lineage reconstruction to characterize the pattern of error introduction

The pattern of errors introduced into the barcode contents was investigated by constructing lineages for individual clusters. We hypothesized that the origin UID would have the most paired-UIDs because the first-copied strand has the highest probability of forming a daughter strand during the entire PCR cycle. We then repeatedly arranged the linked UIDs to complete the path (**Supplementary Fig. 4**). The first-copied strand was defined as the ancestor, with the subsequently produced molecules with UID-pairs defined as descendants. We initially examined whether errors are conserved along generations. To elucidate the error pattern, we focused on clusters containing barcode content in which one or two mismatches were introduced at a not negligible frequency (at least one cluster for the barcode content has a specificity of <90%). From all barcode contents in which an error was observed, we randomly selected 23 barcode contents and constructed a phylogenetic tree (**Fig. 2k**, **Supplementary Fig. 5**).

The experimental topology appeared to be similar to that of the theoretical tree as it was observed that the number of branches toward descendant molecules gradually decreased as generations increased. This trend was reasonable, because the number of daughter strands synthesized can match the number of residual cycles. We concluded that the difference in number of branches and taxa between the theoretical and experimental trees was due to incomplete amplification and loss of molecules during purification.

We then investigated how the errors were introduced. We assumed that errors could be introduced in three stages: (i) the six cycles of amplification for UID-tagging (i.e., polymerase errors); (ii) secondary amplification to attach the sequencing adaptor (i.e., polymerase errors); and (iii) during sequencing (i.e., sequencing errors). We hypothesized that errors introduced in the first stage would be conserved along generations at a high frequency, whereas errors introduced in the second and third stages would produce a pattern of low-rate sporadic errors.

Experimentally, the frequency of errors in individual branch points was low (**Supplementary Fig. 6a**), and few of the errors were conserved along generations (**Supplementary Fig. 6b**). This indicated that most of the observed errors were introduced during secondary amplification or sequencing (i.e., second and third stages). We expected that polymerase errors would be rare with the high-fidelity polymerase: A total of 2,788 molecules were identified experimentally using 88,982 daughter strands, corresponding to 1,067,784 bases for barcode contents (i.e., 12 bases of barcode content per strand × the number of daughter strands). However, the manufacturer’s reported error rate of the polymerase used in this experiment is one error per 3.6 × 10^6^ bases.

Similar patterns were observed in experiments using different polymerases. We chose QIAGEN Multiplex PCR polymerase (designated “QM”), which is derived from family A polymerases and known to have higher error rate than KAPA polymerase, and Phusion polymerase (designated “PH”), which is derived from family B polymerases such as KAPA ^40,41^. A total of 3,488 molecules were obtained using 138,857 daughter strands with QM, and 2,500 molecules were obtained using 96,023 daughter strands with PH (**Supplementary Fig. 7**). Both polymerases exhibited specificity comparable to that of KAPA polymerase, and the specificity increased after correction of one or two base errors introduced into the barcode contents. Polymerase errors were rare in both experiments, even when using QM, which has a higher error rate than family B polymerases (**Supplementary Figs. 8, 9**).

Several advantages of our SPIDER-seq clustering strategy versus conventional methods were noted during clustering and reconstruction of the lineage tree: (i) a greater number of reads is used to remove background errors in amplicon sequencing, (ii) more than one consensus can be created from one molecule to detect extremely low amounts of mutant DNA, and (iii) information regarding error patterns is generated. Analyses of the error pattern provided a guide for the use of SPIDER-seq for identifying ctDNA. Considering that 50,000–90,000 error-reduced consensus sequences were obtained from thousands of initial molecules (**Supplementary Table 5**), we calculated that at a minimum, dozens of clusters would clearly reflect the rare proportion of ctDNA, even if only one or two ctDNA molecules were present among the thousands of initial molecules.

To validate our experimental design, we computationally generated five replicate datasets and reproduced the experiment through simulation. Starting with 1,000 initial molecules, we constructed a library that produced 114,000 virtually generated reads. Excluding instances where polymerase errors (with a high error rate of approximately 10^−5^) occurred during initial UID attachment, only 0.01% of consensus sequences mismatched the original sequence (**Supplementary Fig. 10**). Additionally, in simulations mimicking partial sequencing of the NGS library, we observed similar outcomes to Supplementary Fig. 3 and Fig. 2h, where clusters broke apart, reducing the number of pairs per cluster. Despite this cluster breakage, we recovered approximately 5,000 clusters from the starting 1,000 initial molecules which can improve sensitivity. This demonstrates that our method can robustly perform error correction, even in the presence of molecule loss during sequencing and experiments, as long as a high-fidelity polymerase (with an error rate below 10^−6^, commonly used in NGS workflows) is employed.

### Detection of mutations with an allele frequency of 0.125%

To confirm the utility of SPIDER-seq for ctDNA detection, we tested mock cfDNA references containing mutations in dozens of genes with the average variant allele frequency adjusted to 1, 0.5, 0.125, and 0% (i.e., control). We particularly targeted the *BRAF* gene harboring the p.V600E mutation and amplified the *BRAF* V600 locus using UID-containing primers and eight amplification cycles. Using only 12.2–15.8 ng (3,697–4,788 hGEs) of mock cfDNA, an average of 215,551 daughter strands were obtained, and an average of 113,234 clusters were created using the strands via construction of the peer-to-peer network. Among the clusters, an average of 42,795 consensus reads with ≥2 supporting UID-pairs were obtained. From the consensus reads, mutations could be detected at a variant allele frequency of 0.125%, and changes in other bases were rare (**Fig. 3a, Supplementary Fig. 11, and Supplementary Table 6**). To compare method performance in terms of variant detection, the same data were analyzed by simple base-counting or creation of a consensus read for each UID-pair (i.e., without constructing a cluster) (**Fig. 3b**). The results showed that the UID-based consensus method significantly reduced the error rate compared to the raw data (P = 4.88 × 10^−4^, Wilcoxon signed rank test). Furthermore, the cluster-based consensus method demonstrated an even greater reduction in error rate compared to both the raw data and the UID-based consensus (P = 4.88 × 10^−4^ between raw data and SPIDER-seq, and P = 4.88 × 10^−4^ between UID-based consensus and SPIDER-seq, Wilcoxon signed rank test).

From tens to hundreds of consensus reads representing the p.V600E mutation were identified in the 0.125% variant allele frequency sample (**Supplementary Table 6**). We expected that there would be ≤10,000 starting strands for amplification (i.e., 2 strands × 5,000 haploids); therefore, the ideal number of mutant strands should be ≤12. These data indicate that redundant reads obtained using our SPIDER-seq method provide a buffer against the loss of molecules during preparation. This buffering function is expected to be particularly beneficial in the monitoring of known variants.

We also investigated the errors introduced in target loci. In addition to the p.V600E mutation (corresponding to a genomic A to T mutation), a rare fraction of A to G mutations was observed in two replicates of the 1% variant allele frequency mock cfDNA samples, and one A to T mutation was observed in one replicate of the control sample (**Supplementary Table 6**). Reconstruction of the lineage of the clusters showed that the unexpected base changes were highly conserved among all branch points, suggesting the occurrence of polymerase errors (**Supplementary Fig. 12**). We concluded that these errors were introduced during the early cycles of amplification, as the proportion of errors was 100% in all branch points in the clusters and the errors were not observed in all replicates. Compared to the conditions of the model experiment above, the possibility of having polymerase errors is higher because two more cycles were added in the amplification step, and a greater number of daughter strands were sequenced. Similarly, a high frequency of errors was observed in some clusters of the locus near the mutation (**Supplementary Table 7**), and these errors were observed in particular replicates, similar to the case of the locus in which the mutation was detected. These results suggest that polishing by comparing mutant information for multiple replicates and control samples would be helpful in clarifying whether an observed base change is an error or a mutation.

To determine the minimal data requirement for analyzing low-frequency mutations, sequencing reads were down-sampled to a depth of 10,000–10,000,000 reads (**Materials and Methods**). We found that 100,000 reads were sufficient to detect a mutation at a variant allele frequency of 0.125% (**Supplementary Fig. 13a**). This result indicates that mutations can be identified in several loci using a MiniSeq Rapid Kit, which can generate 2 Gb of data within 5 hours. This approach would therefore be advantageous when examining a small number of individual samples at irregular intervals, as in the case of MRD monitoring. However, to obtain consensus using a larger number of daughter strands, we expect that >100,000 reads would be required (**Supplementary Fig. 13b**).

### Multiplex detection of mutations in 10 genes

Next, we demonstrated that SPIDER-seq method can be extended to simultaneously detect mutations in multiple locations. For this purpose, we employed a multiplex PCR approach that uses a mixture of primers in a single amplification reaction with QM polymerase, which is widely used for specific and sensitive multiplex PCR analyses. A total of nine substitution mutations and one deletion mutation (EGFR p.E746_A750del) were selected among the mutations in the mock cfDNA (**Supplementary Table 8**). The mock cfDNAs, with the average variant allele frequency adjusted to 0.25, 0.125, or 0%, were then amplified and analysis of the mutations was performed. The variant allele frequencies of the targeted substitution mutations agreed well with the known variant allele frequencies provided by manufacturer. The average reduced error rate was 0.02369%, which was higher than the experimentally targeted single *BRAF* mutation position (0.002628%) (**Fig. 3b–e and Supplementary Fig. 14**). We concluded that the QM polymerase introduced more errors than KP polymerase during eight amplification cycles ^40^.

To examine indel mutations, we developed an algorithm distinct from that used for substitution calling. Whereas substitutions can be examined by counting A, T, C, and G bases in a given locus, indel mutations cannot be examined in the same manner. As the size of an insertion or deletion can vary in a given locus, indel mutations cannot be counted before they are specified. Therefore, we devised the following three-step strategy: (i) generate a variant call format (vcf) file using an indel calling program such as VarDict ^42^ or list known indels to be queried; (ii) construct a peer-to-peer network to define clusters; and (iii) check the consistency of indels from the reads of each cluster. For this experiment, we generated a vcf file by listing the known deletion mutation in the mock cfDNA sample. As a result, we found the consensus reads representing the deletion with the expected deletion frequency (**Fig. 3c**). We conclude that false-positive indel identifications would be rare, because a 15-nt deletion error would rarely occur naturally during preparation and sequencing steps. Therefore, the highly specific identification of the deletion mutation indicated that the genotype of the original molecule was retained during cluster formation. Taken together, these data demonstrate that SPIDER-seq can be used to identify both substitution and indel mutations in ctDNA within multiple targeted loci in a single experiment.

### Use of an alternative library for hybridization capture

Although SPIDER-seq method was designed to reduce errors in a small number of loci based on an amplicon sequencing protocol, the method also uniquely enables reconstruction of phylogenetic trees to track error patterns. Therefore, in tandem with the purpose of reducing errors, we adjusted the method so that it could be used alternatively with a library prepared based on an adaptor ligation protocol. We hypothesized that this would make it possible to investigate the most error-prone step during target sequencing via hybridization capture methods. For this purpose, we modified our strategy for the amplification step for preparation of a shot-gun sequencing library based on an adaptor ligation protocol. We selected three bases as a sample barcode and five bases as a UID sequence and utilized them to amplify an adaptor-ligated product to be sequenced as part of the “index read” (**Fig. 4a**), while the standard ligation-based protocol for hybridization capture uses adaptors containing UIDs themselves^15^. Using the shot-gun sequencing library, a genomic coordinate was used as a secondary identifier to supplement the low diversity of the five-base UID. To test the feasibility of constructing a peer-to-peer network from a shot-gun DNA library, we prepared libraries from mock cfDNA in which the average variant allele frequency was adjusted to 0, 0.125, 0.25, 0.5, or 1% using eight amplification cycles for UID-tagging. A hybridization capture experiment was then performed using a panel targeting 68 genes covering 24 substitution mutations and 4 non-homopolymer indel mutations in the mock cfDNA (**Supplementary Table 9**). As a result, we obtained 355,921× coverage on average per-target mutations (**Supplementary Table 10**) and 21 substitution and 4 indel mutations, with coverage >100,000, corresponding to the minimal depth for detecting a mutation at a variant allele frequency of 0.125%, were subjected to construct the peer-to-peer network.

**Fig. 4.**
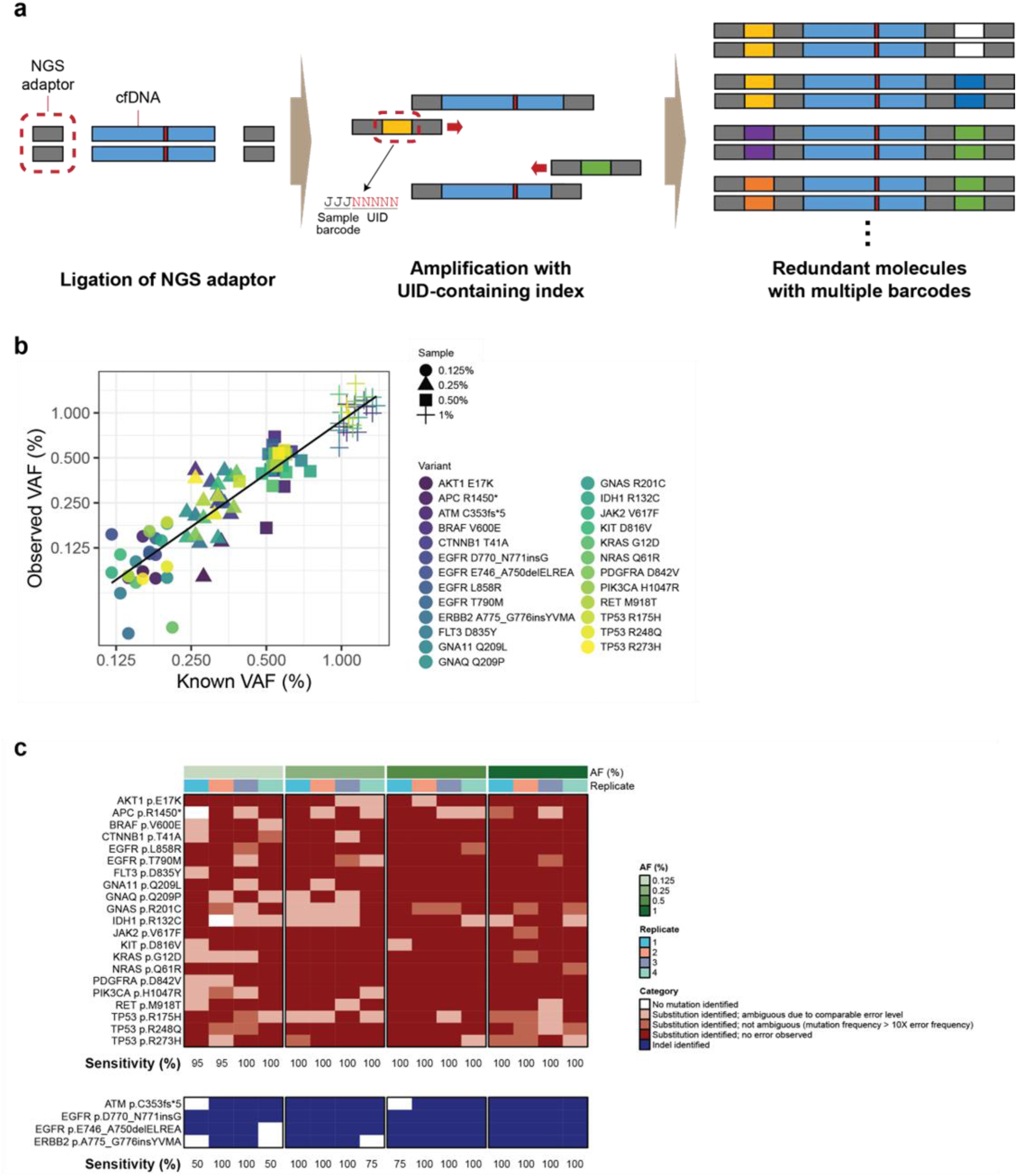
Application of the SPIDER-seq method to the library prepared using adaptor ligation. (**a**) Schematic illustration of CID-based barcoding for preparation of the shot-gun sequencing library. (**b**) Comparison of VAFs observed using SPIDER-seq with known VAFs provided by the manufacturer. For each sample and variant, the mean observed VAF from replicate experiments is presented. The line is a linear fit. Pearson *r* = 0.93944. (**c**) Identification of mutations from hybridization capture data at AF values of 1, 0.5, 0.25, and 0.125%. Each column corresponds to a single sample in a single replicate experiment.

To construct the peer-to-peer network, only those UIDs having the same genomic coordinates were used. On average, 6,872 clusters were observed from 25 loci (**Supplementary Table 11**), and the size of the clusters varied (**Supplementary Fig. 15a**). Based on the consensus reads obtained from the clusters, the mutant allele frequencies of both substitution and indel mutations exhibited high concordance with the expected variant allele frequencies provided by the manufacturer (**Fig. 4b**). In addition, the error rate declined by 5.453-fold by using the CID-based consensus over simple base-counting from raw bam. This result demonstrates that SPIDER-seq can be adapted to ligation-based protocols. However, there was a non-negligible difference in performance compared to the amplicon sequencing protocol. First, the sensitivity was not 100% in the 0.5, 0.25, and 0.125% variant allele frequency samples (**Fig. 4c**). This decrease in sensitivity was presumably caused by a loss of molecules during the additional experimental steps of hybrid capture compared to the amplicon sequencing protocol. Second, the basal level of errors observed in the data without creating consensus (i.e., 0.0672%) was higher compared to direct PCR amplification of the *BRAF* locus (i.e., 0.0202%), even though KP polymerase was used in both experiments (**Fig. 3b and Supplementary Figs. 15b, c**). Third, CID-based consensus showed only a small, though statistically significant, improvement compared to UID. We concluded that a larger amount of starting material and more sequencing data would be required to enhance the sensitivity. In addition, more-stringent criteria would be needed to filter out false-positive results presumably resulting from the higher error rate compared to that of the amplification-based preparation. Nonetheless, application of SPIDER-seq resulted in a marked reduction in the error rate compared with analysis of raw data.

We hypothesized that errors could be introduced during four steps: (i) preparation of the pre-capture library (i.e., polymerase errors). In this case, errors would be conserved in descendant molecules at high frequency. (ii) During hybridization capture due to oxidative damage ^43^. Errors introduced in this step would be focused on a particular node at a high frequency, but they would not be conserved. (iii) During post-capture amplification (i.e., polymerase errors). (iv) During sequencing (i.e., sequencing errors). Errors introduced through steps (iii) or (iv) would be sporadic and exhibit a low frequency. To visualize these error patterns, we reconstructed phylogenetic trees of clusters exhibiting a non-reference genotype (**Supplementary Figs. 16–19**). Most of the errors appeared to be conserved across the entire branch, indicating errors introduced during step (i). However, most clusters of the non-reference genotype were comprised of only two daughter strands, making it difficult to define the most error-prone step. We hypothesized that case (ii) could produce a similar pattern when error-containing clusters were split into small clusters due to experimental loss of molecules.

Taken together, these data demonstrate that our SPIDER-seq approach can be applied to ligation-based protocols, with sufficient sensitivity to detect mutations at a 0.125% variant allele frequency. However, due to loss of molecules, the sensitivity is somewhat low and the error rate is high compared to amplification-based protocols. Thus, an amplification-based method would be a better option in terms of ctDNA loss than a hybridization-based method when starting with a low number of molecules.

## Discussion

In this study, we developed an alternative method for accurate detection of variants present at a low allele frequency. With previously developed methods, large branches are based on hybridization capture, which can be time consuming (requiring 2 days), and these methods require the synthesis of baits prior to the primary experimental steps. Other methods are based on PCR with a limited number of amplification cycles, which can result in loss of molecules or an increased number of enzymatic reactions after the first round of PCR ^23,30–32^. In contrast, SPIDER-seq enables the detection of a mutant allele at a variant allele frequency of 0.125–1% via just two rounds of general PCR. Considering its simplicity and versatility, SPIDER-seq is suitable for use in diagnosing cancer relapse in local institutions where labor and equipment are limited.

The novelty of SPIDER-seq is its use of peer-to-peer networks for error reduction. In the ligation-based method, daughter strands originating from one starting molecule have identical UIDs, so the daughter strands can be used to create a consensus sequence of the starting molecule. However, tagging daughter strands with identical UIDs is difficult with PCR because each cycle results in multiple combinations of daughter strands by overwriting. An advantage of SPIDER-seq is that it enables the creation of a new identifier by joining combinations generated during PCR through a peer-to-peer network. As we described in the **Results** section, construction of a peer-to-peer network enables the collection of more reads compared with the collection of reads based on UID-pair information. Moreover, molecules can be rearranged in the order in which they were generated, similar to a phylogenetic tree. In this study, phylogenetic tree reconstruction was conducted only for tracking error-introduced branch points, but we expect that our approach could be expanded to other experiments, such as lineage tracing. Current methods for lineage tracing involve the analysis of barcode sequences edited using CRISPR ^44,45^ or Cre-*loxP* ^46^. As alterations in the sequences are usually permanent, lineage barcoding after all editable sites have been altered is not cumulative. This means that tracing is not possible in a long-term experiment. A system in which two barcode sequences are alternately overwritten via homology-directed repair could make it possible to connect very long generations by linking peer-to-peer networks.

Although we demonstrated the novelty and feasibility of SPIDER-seq, further work is needed to reduce the broken clusters in order to more accurately reflect molecular information during amplification. Broken clusters can result in a particular molecule being over-counted, leading to potential allele bias. Broken clusters can also produce numerous small clusters containing only one UID-pair, which could hamper error correction due to an insufficient number of reads for creating consensus. According to our simulation data (Supplementary Fig. 10), broken clusters primarily occur because of molecule loss during the sequencing library preparation and because only a portion of the sample is taken for sequencing. Therefore, optimizing the experiment protocol to minimize sample loss is crucial. Additionally, errors in the UIDs can also cause breakage. When errors occur in both UIDs, even if a molecule is not dropped out, it may appear disconnected from other reads in the original cluster. Using high-fidelity polymerase (with an error rate of 1 per 3.6 × 10^6^ bases commonly used in NGS library preparation) can help mitigate this issue to some extent, and adopting an algorithm that corrects errors in UIDs can improve the overall integrity of the sequencing results. By incorporating such improvements, we believe that SPIDER-seq would be an ideal option for deep sequencing to detect mutations in low-frequency alleles.

## Data availability

Raw sequence data are available in Sequence Read Archive (SRA) under project number PRJNA693080.

## Code availability

The code supporting our analysis is available at https://github.com/limhs/SPIDERseq.git.

## Acknowledgments

This work was supported by the National Research Foundation of Korea (NRF) grant funded by the Korea government (MSIT) (No: 2021R1A2C2008490) and by the Korean Government (RS-2024-00338316).

## Author contributions

D.B. and J.H.L. supervised the study. H.L., S.J. and T.K. designed and performed the experiments, analyzed the data, and wrote the manuscript.

