## Supplementary Data for "Correcting errors in PCR-derived libraries for rare allele detection by reconstructing parental and daughter strand information"

### Supplementary Materials

#### Supplementary Note

##### Total number of molecules obtained from a single-stranded starting molecule after a given number of cycles

Assuming that the polymerase catalyzes the synthesis of molecules with 100% efficiency in each cycle, the number of molecules can be considered to double every cycle. Thus, the total number of molecules obtained from a starting molecule would be $2^n$ after n-th cycle. Molecules lacking a left or right UID cannot be sequenced. Therefore, starting molecules lacking a UID on 5’ side (see **Figure 1B and Supplementary Figure S1**) are not sequenced. Thus, the total number of molecules that could be subjected to sequencing is $2^n - 1$. Subtracting a value of ‘1’ represents exclusion of the starting molecule.

##### Number of daughter strands per cluster

We assumed that first-copied strands would be generated in each cycle and that the number of molecules per cluster could be estimated by defining the first-copied strand as the starting molecule. Assuming that the first-copied strand was generated in the i-th cycle, the number of remaining cycles is $n - i$. The number of molecules derived from the first-copied strand would then be $2^{n-i}$ ($i = 1$ to $n$).

##### Number of UIDs

Among the newly generated molecules in a given cycle, half of the molecules can be considered to have been generated by attaching a new left UID and the other half as being generated by attaching a new right UID. Therefore, among $2^{n-i}$ molecules in the cluster that originated from the first-copied strand in the i-th cycle, we can assume that $2^{n-i-1}$ molecules (half of the molecules in the cluster) were generated by attaching a new left UID and that the remaining $2^{n-i-1}$ molecules were generated by attaching a new right UID. Therefore, the largest possible number of UIDs per cluster is $2^{n-2}$ for clusters that originated from the first-copied strand in the first cycle ($i = 1$).

#### Supplementary Figures

**Fig. S1**

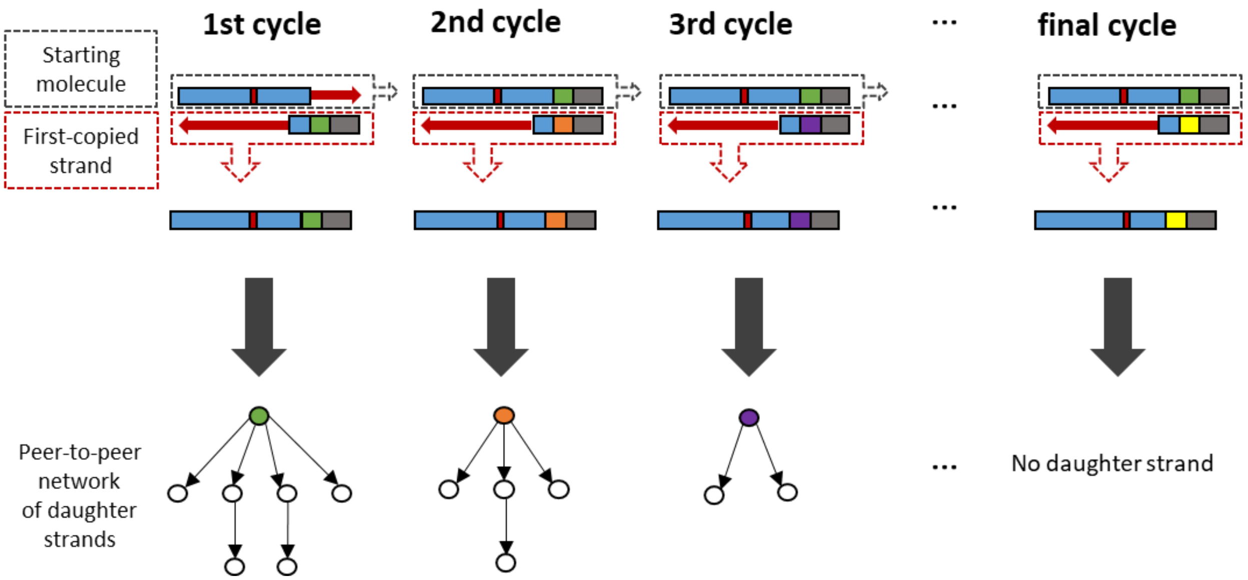


**Supplementary Figure 1.** Schematic illustration describing the process of triggering multiple networks using one starting molecule.

**Fig. S2**

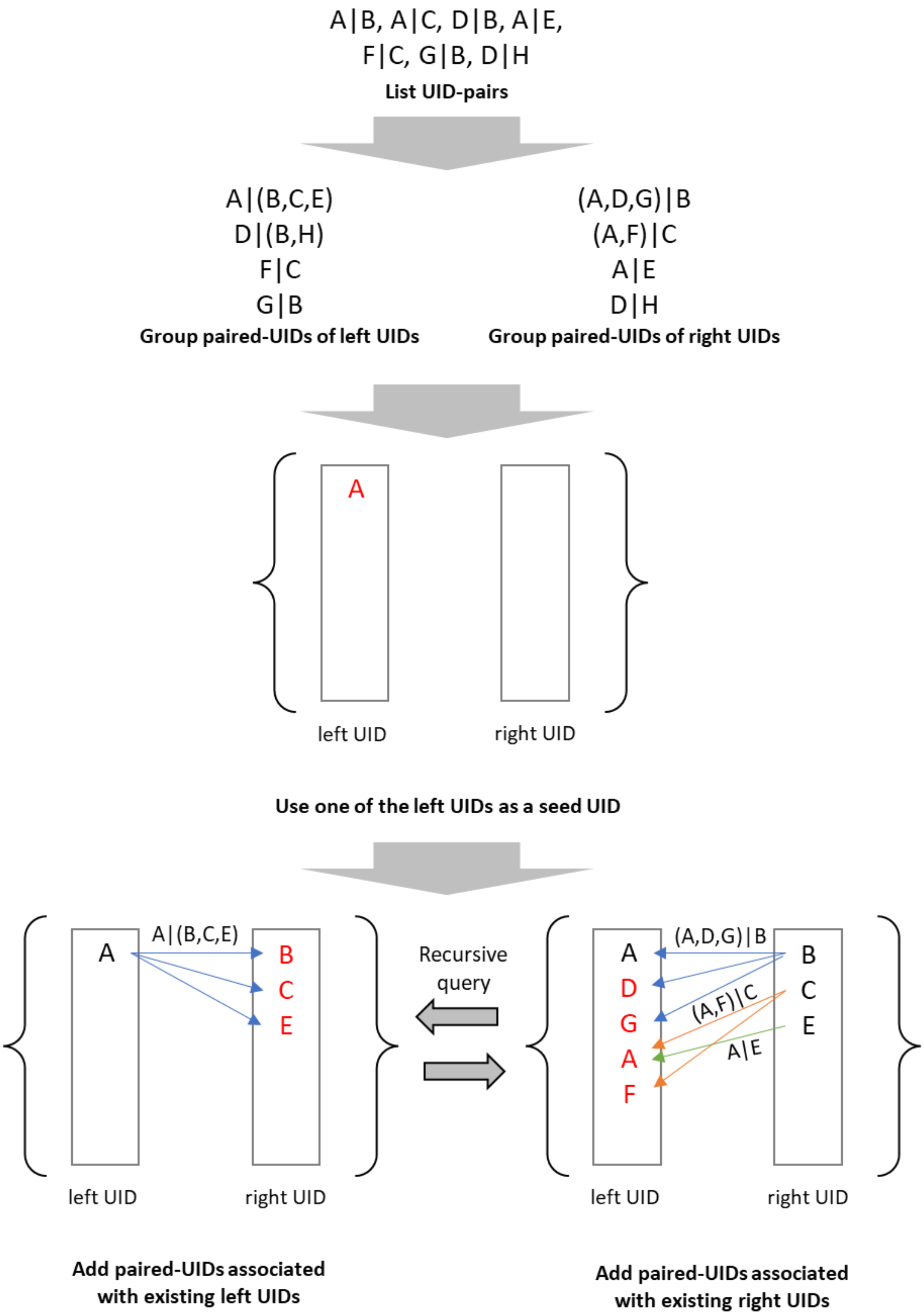


**Supplementary Figure 2.** Schematic illustration of the algorithm for linking UIDs. Paired-UIDs associated with existing UIDs are recursively added until no paired-UIDs remain. Red font indicates newly added UIDs.

**Fig. S3**

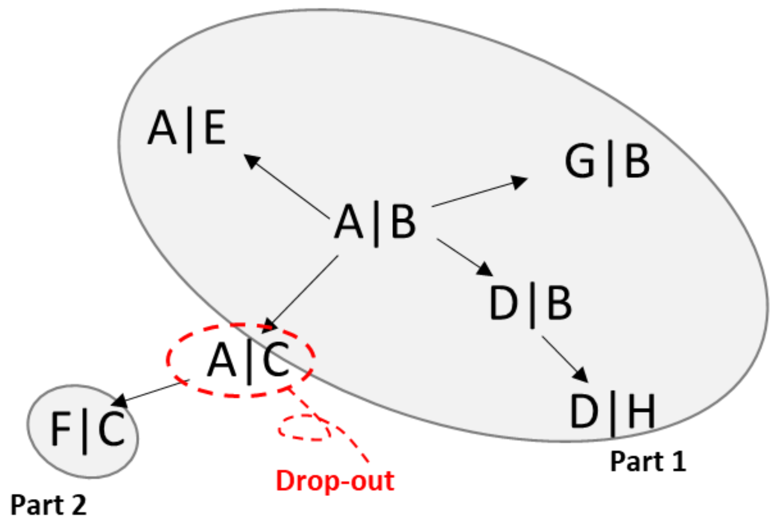


**Supplementary Figure 3.** Illustration describing the case of a broken cluster. Loss of the UID-pair in the middle of the connection separates the cluster into two parts.

**Fig. S4**

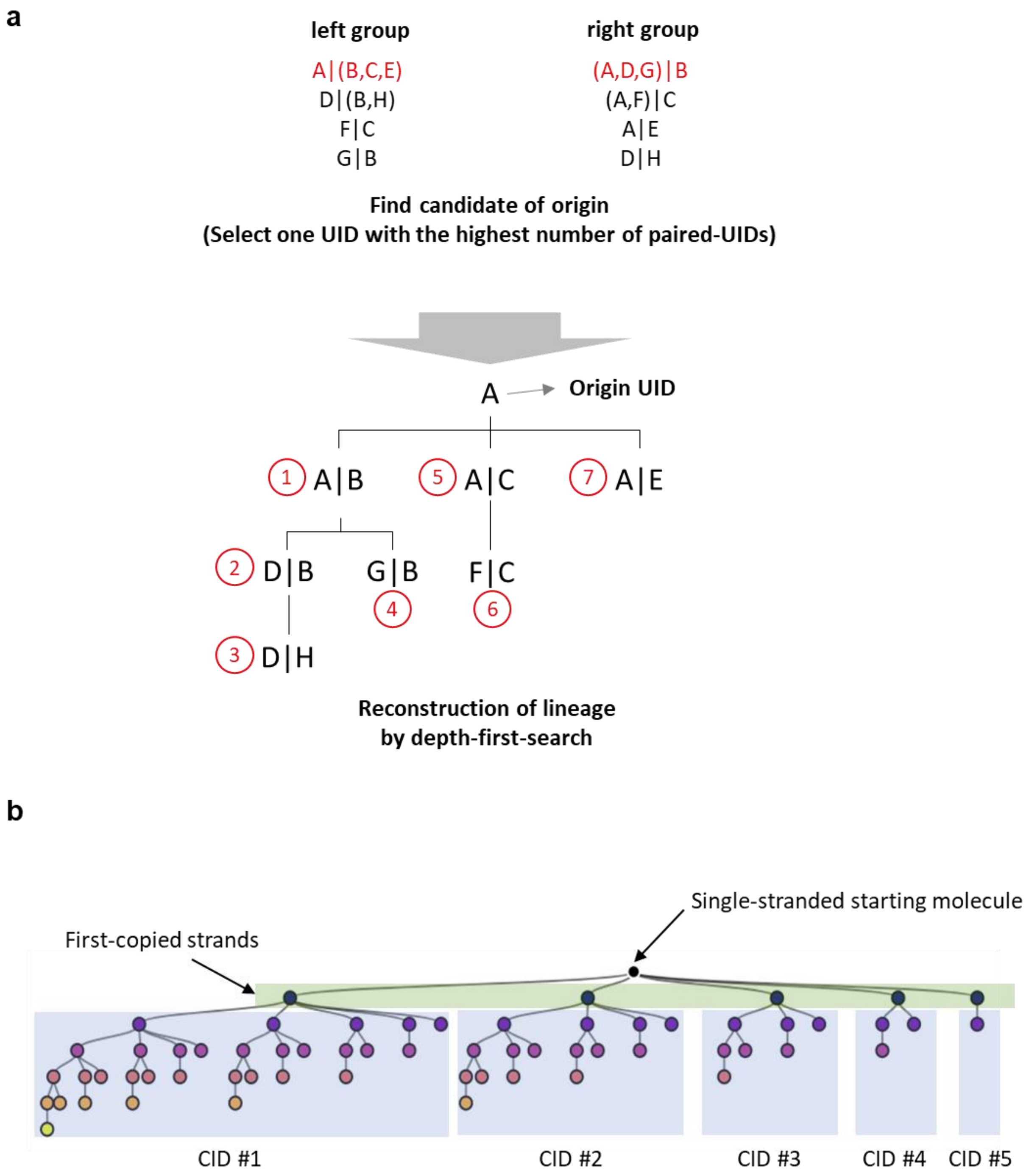


**Supplementary Figure 4.** Illustration of the concept for lineage construction. (**a**) The origin UID is placed at the position of the root, and branches are extended using a depth-first search method. (**b**) Dendrogram of the theoretical lineage of strands produced from a starting molecule in six cycles of amplification. The starting molecule is denoted as a black circle, and the strands produced by amplification are denoted as circles with color gradation of indigo to yellow with black border. Five clusters are generated by amplification and denoted as clades (CID = cluster identifier).

**Fig. S5**

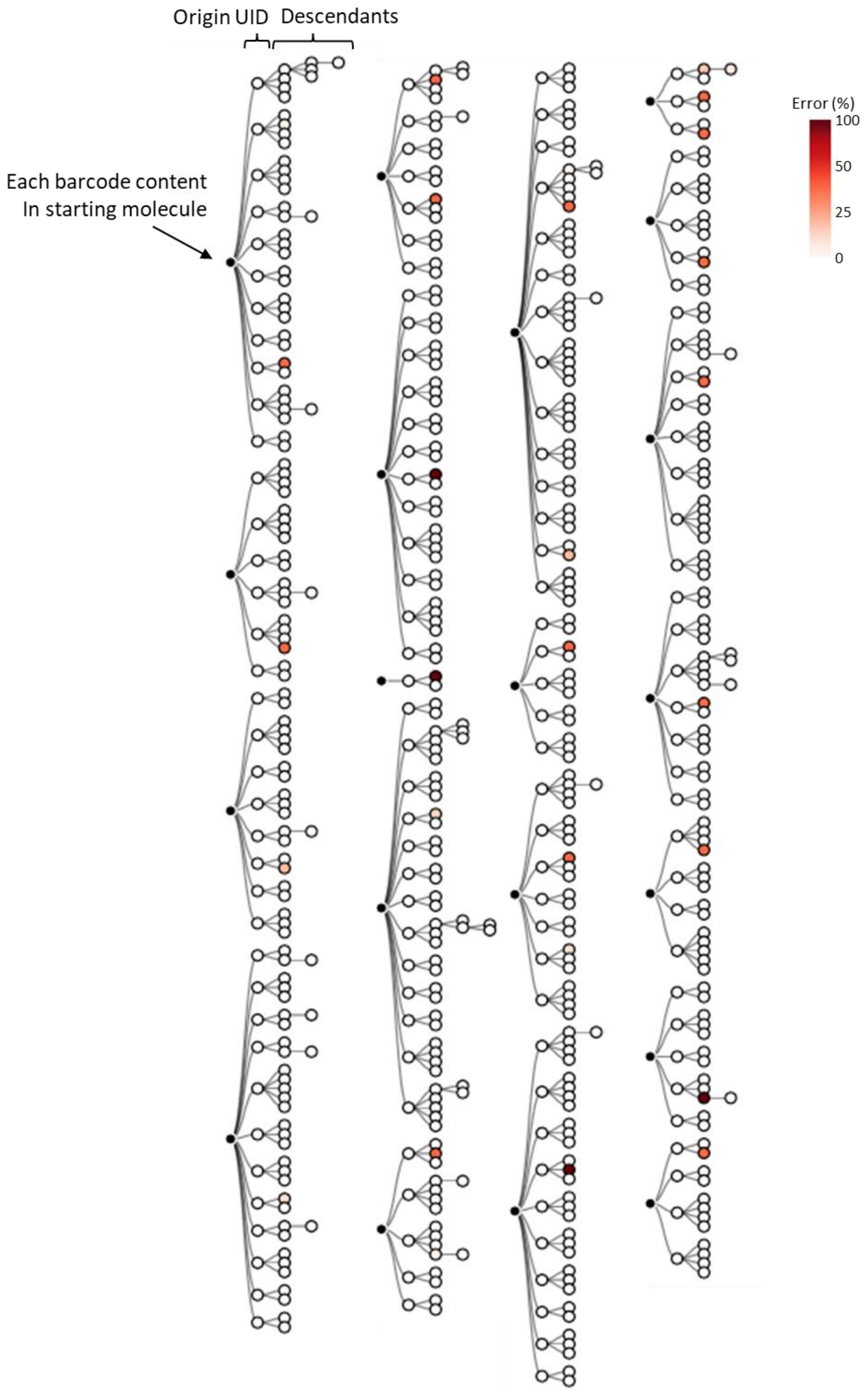


**Supplementary Figure 5.** Phylogenetic tree obtained from clusters with specificity <90%. Twenty barcodes were randomly selected to show the error pattern.

#### Fig. S6

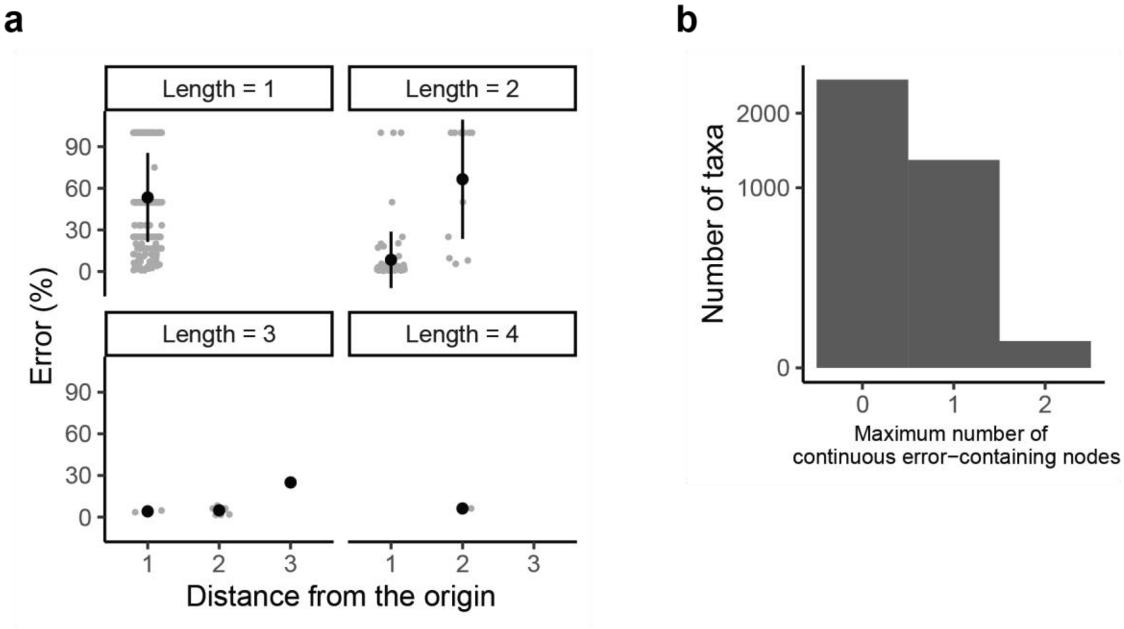


**Supplementary Figure 6.** Analysis of errors introduced into branch points. (**a**) The frequency of errors was low for most taxa. Error rate (%) is plotted in the given length of the branch. Black dots indicate mean values of the error frequency and error bars indicate standard deviations. (**b**) Distribution of the maximum number of continuous error-containing nodes.

**Fig. S7**

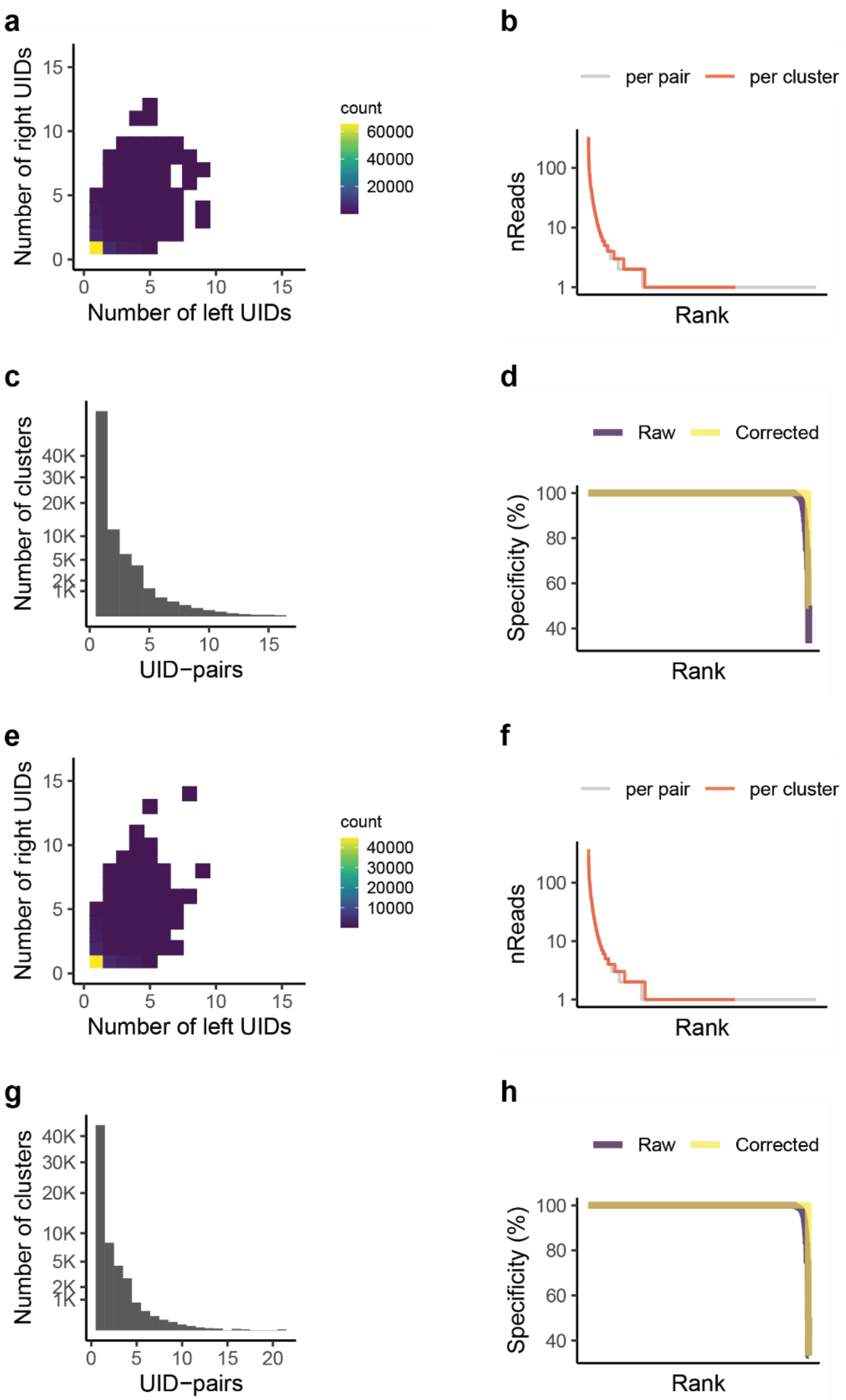


**Supplementary Figure 7.** Analysis of clusters in QM and PH experiments. (**a–d**) Results obtained with QM polymerase. Distribution of cluster size (**a**); the number of reads per UID-pair and per cluster, in which pairs and clusters are given in ranked order (**b**); distribution of UID-pairs per cluster (**c**); and specificity (%) of the clusters before and after barcode content correction within a hamming distance of 2, in which clusters are given in ranked order (**d**). (**e–h**) Results obtained with PH polymerase. Distribution of cluster size (**e**); the number of reads per UID-pair and per cluster, in which pairs and clusters are given in ranked order (**f**); distribution of UID-pairs per cluster (**g**); and specificity (%) of the clusters before and after barcode content correction within a hamming distance of 2, in which clusters are given in ranked order (**h**).

**Fig. S8**

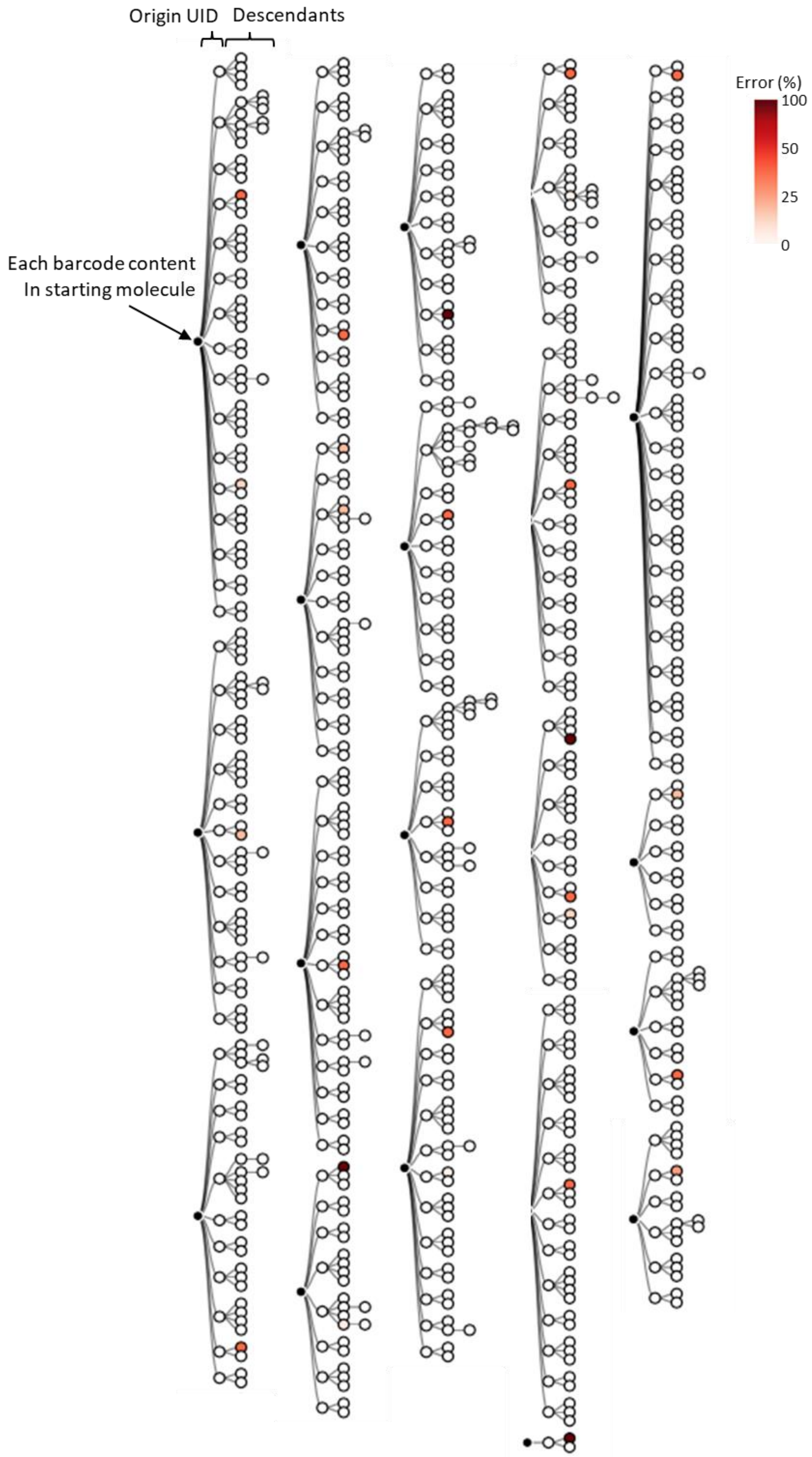


**Supplementary Figure 8.** Phylogenetic tree for QM polymerase obtained from clusters with specificity <90%. Twenty barcodes were randomly selected to show the error pattern.

**Fig. S9**

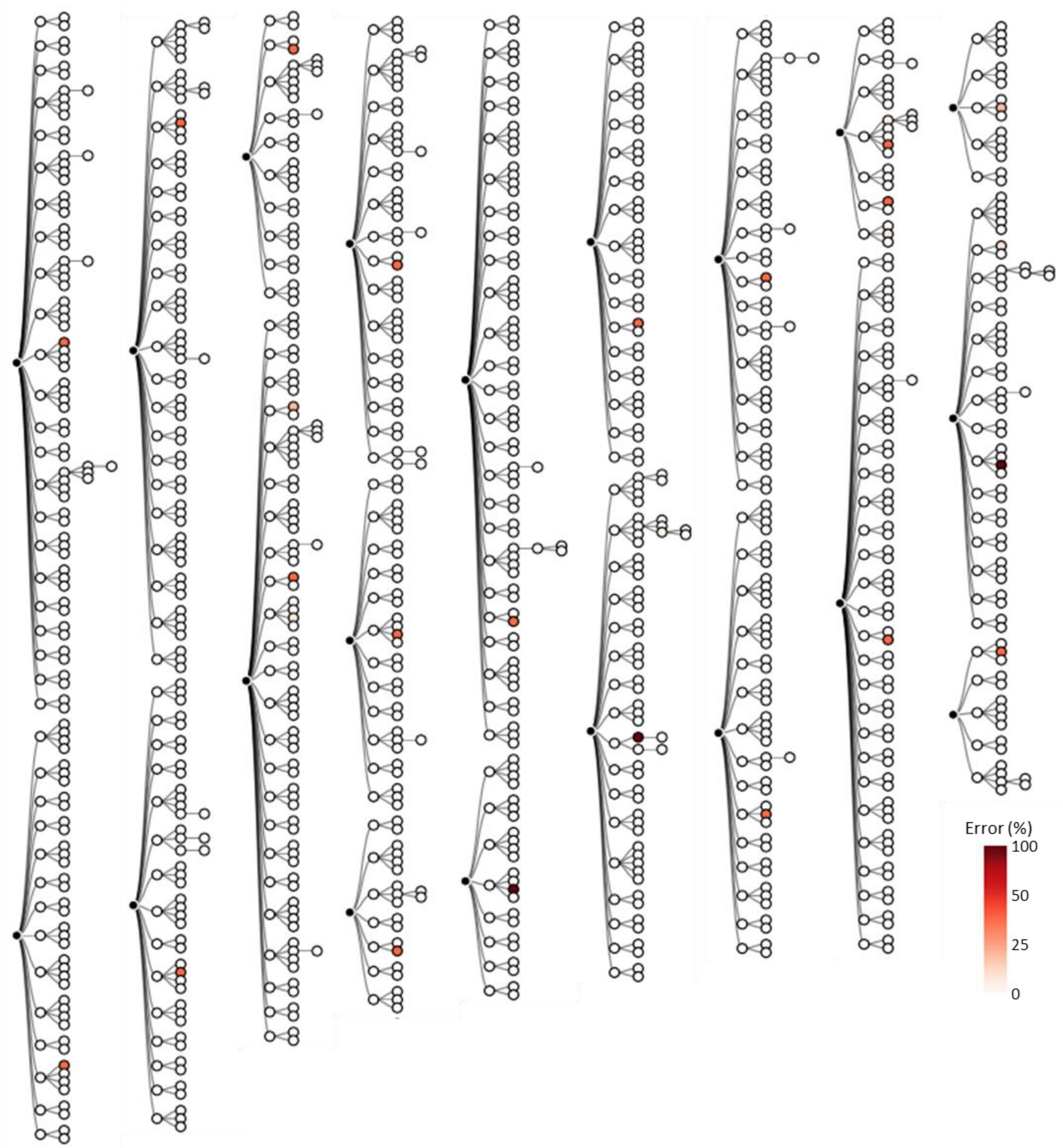


**Supplementary Figure 9.** Phylogenetic tree for PH polymerase obtained from clusters with specificity <90%. Twenty barcodes were randomly selected to show the error pattern.

**Fig. S10**

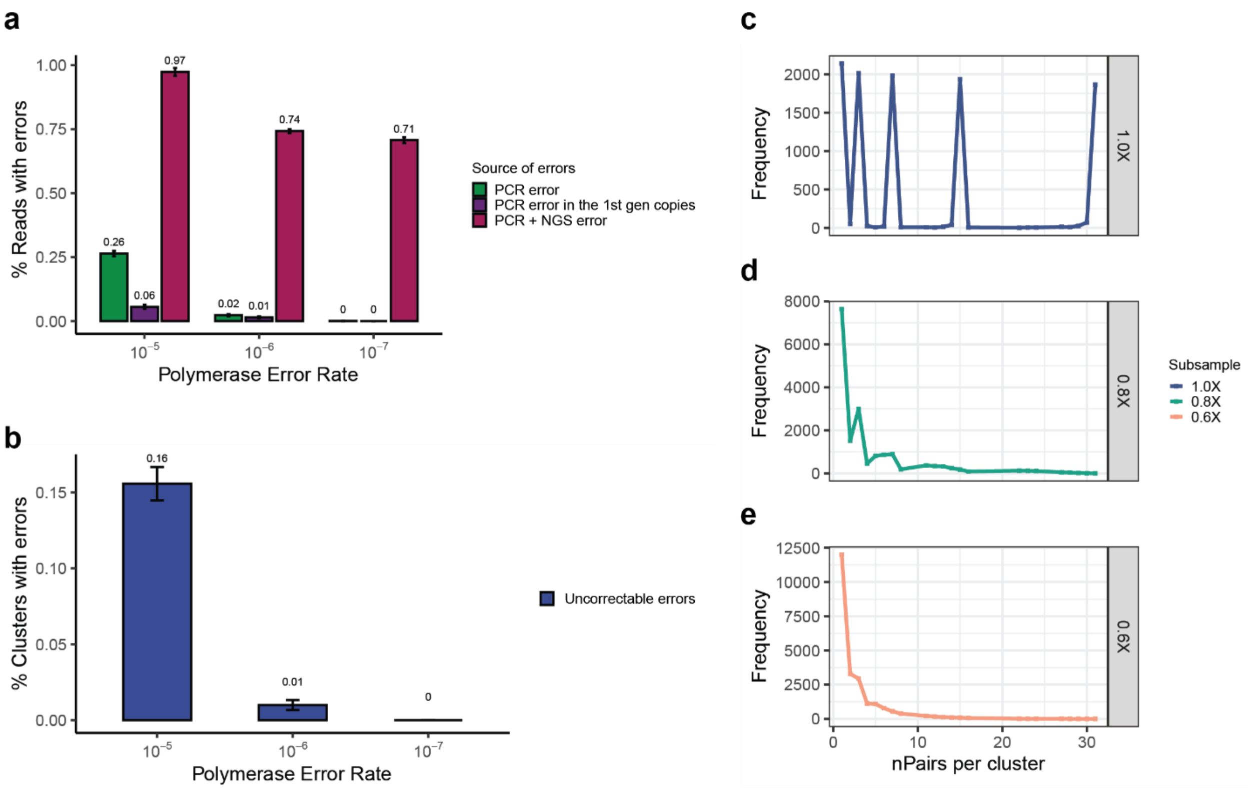


**Supplementary Figure 10.** Computational simulation results. An oligonucleotide-based library was reproduced through computational simulation, incorporating PCR errors (ranging from $10^{-5}$ to $10^{-7}$) and NGS errors (0.1%). **(a)** The major source of error was found to be NGS error, regardless of the polymerase fidelity. PCR errors occurring in the first PCR cycle remained uncorrectable under the condition with the highest error rate ($10^{-5}$) and 0.01% of clusters from $10^{-6}$ **(b)**. **(c–e)** The occurrence of cluster fragmentation due to subsampling, simulating the scenario of using a portion of the NGS library. The original dataset, shown as 1X **(c)**, exhibited multiple peaks, which fragmented into smaller clusters as seen in the subsampled datasets of 0.8X **(d)** and 0.6X **(e)**. Error bars indicate SEM.

**Fig. S11**

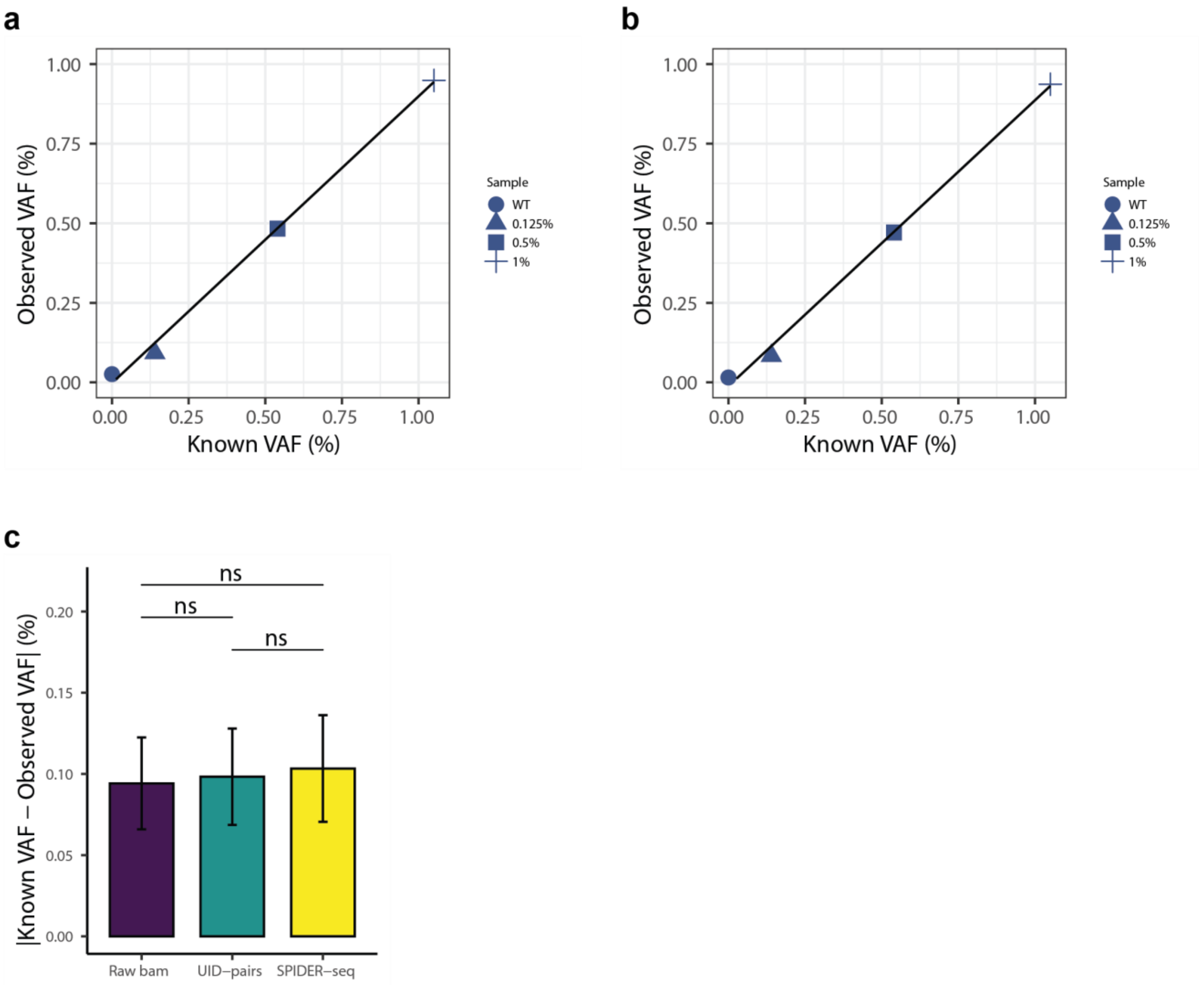


**Supplementary Figure 11.** Known VAF and observed VAF in the BRAF sequencing dataset. (**a**–**b**) Correlation between known VAF and observed VAF in raw data (**a**) and UID-pair-based consensus (**b**). (**c**) Absolute difference between known and observed VAF, demonstrating consistency across error reduction methods ($P$ = 0.266 between raw bam and UID-pairs, $P$ = 0.339 between raw bam and SPIDER-seq, and $P$ = 0.677 between UID-pairs and SPIDER-seq, determined by Wilcoxon signed rank test). Error bars represent the standard error of the mean. ns, not significant.

**Fig. S12**

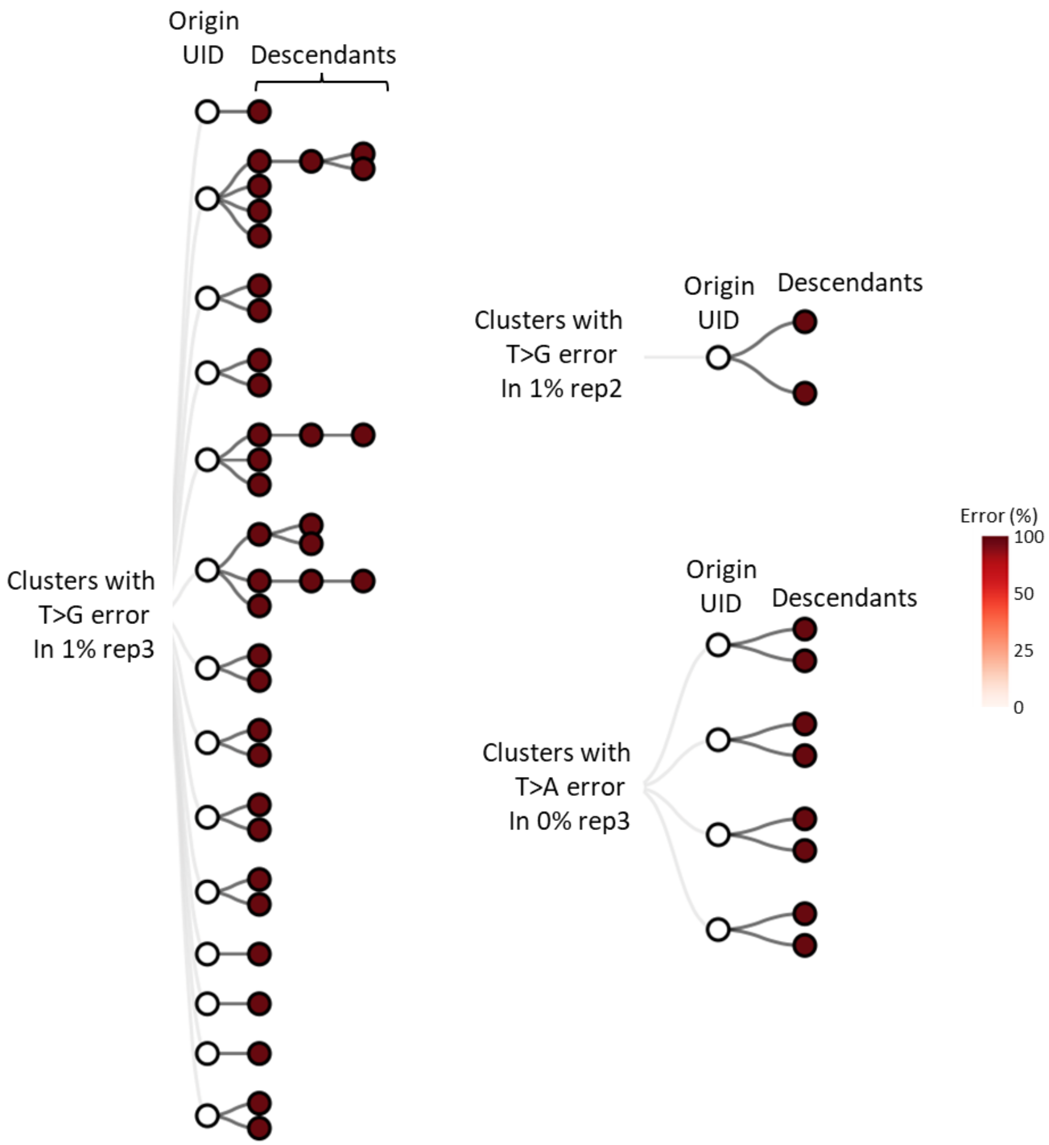


**Supplementary Figure 12.** Phylogenetic tree of clusters exhibiting a non-reference genotype.

**Fig. S13**

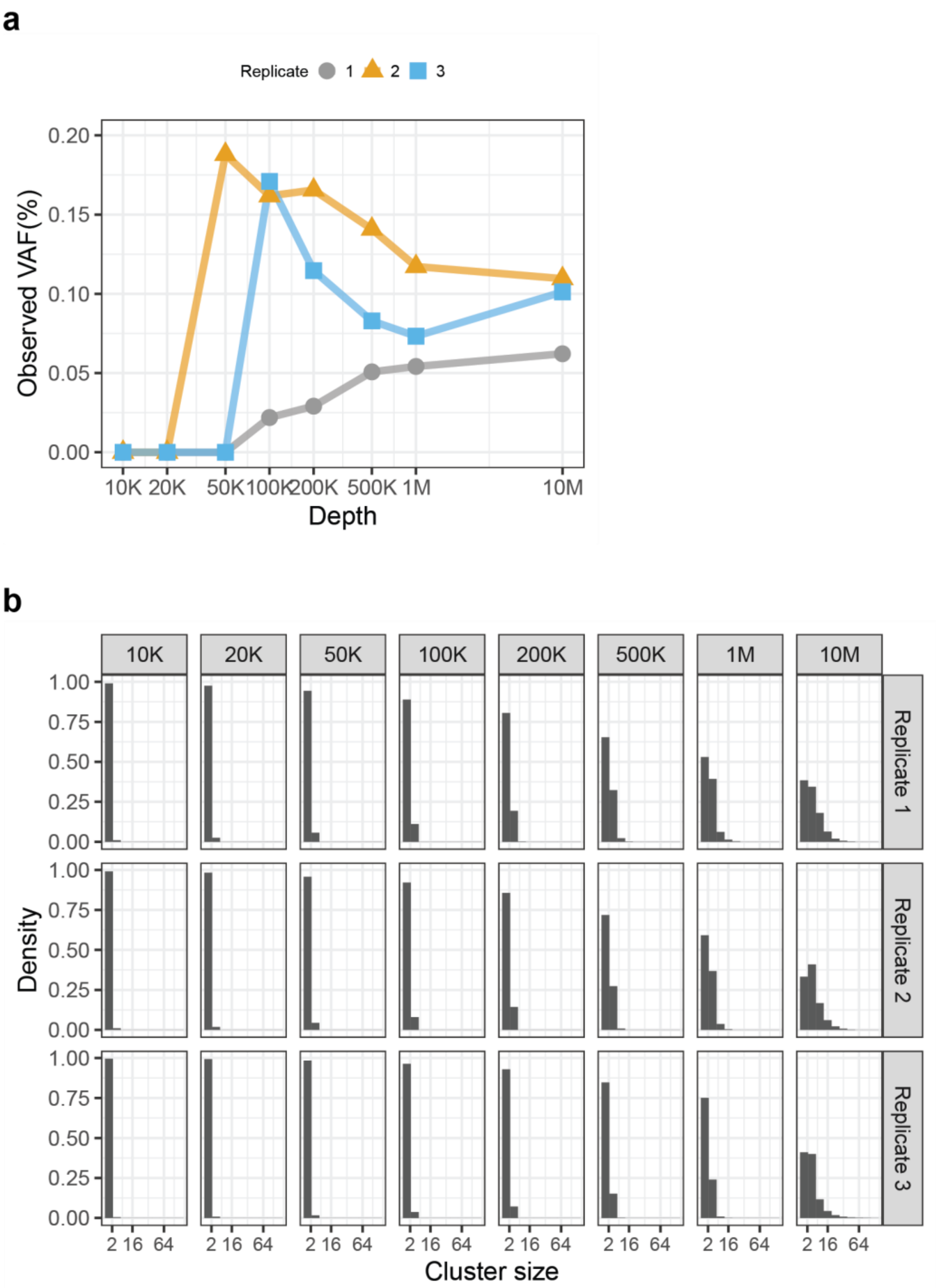


**Supplementary Figure 13.** Minimum data requirement for analyzing 0.125% of mutations. (**a**) VAF (%) observed from down-sampled data. (**b**) Distribution of cluster size for a given depth.

#### Fig. S14

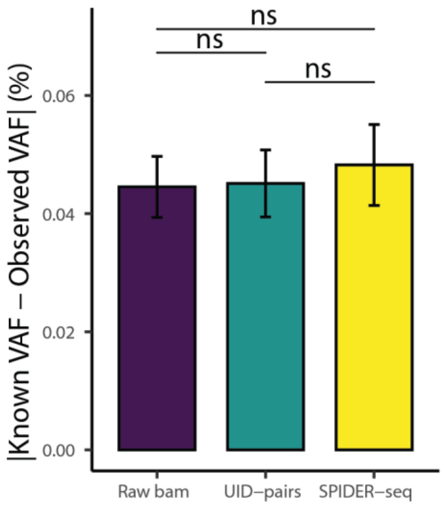


**Supplementary Figure 14.** Absolute difference between known and observed VAF in the 10-gene sequencing dataset (*P* = 0.539 between raw bam and UID-pairs, *P* = 0.286 between raw bam and SPIDER-seq, and *P* = 0.214 between UID-pairs and SPIDER-seq, determined by Wilcoxon signed rank test). Error bars represent the standard error of the mean. ns, not significant.

**Fig. S15**

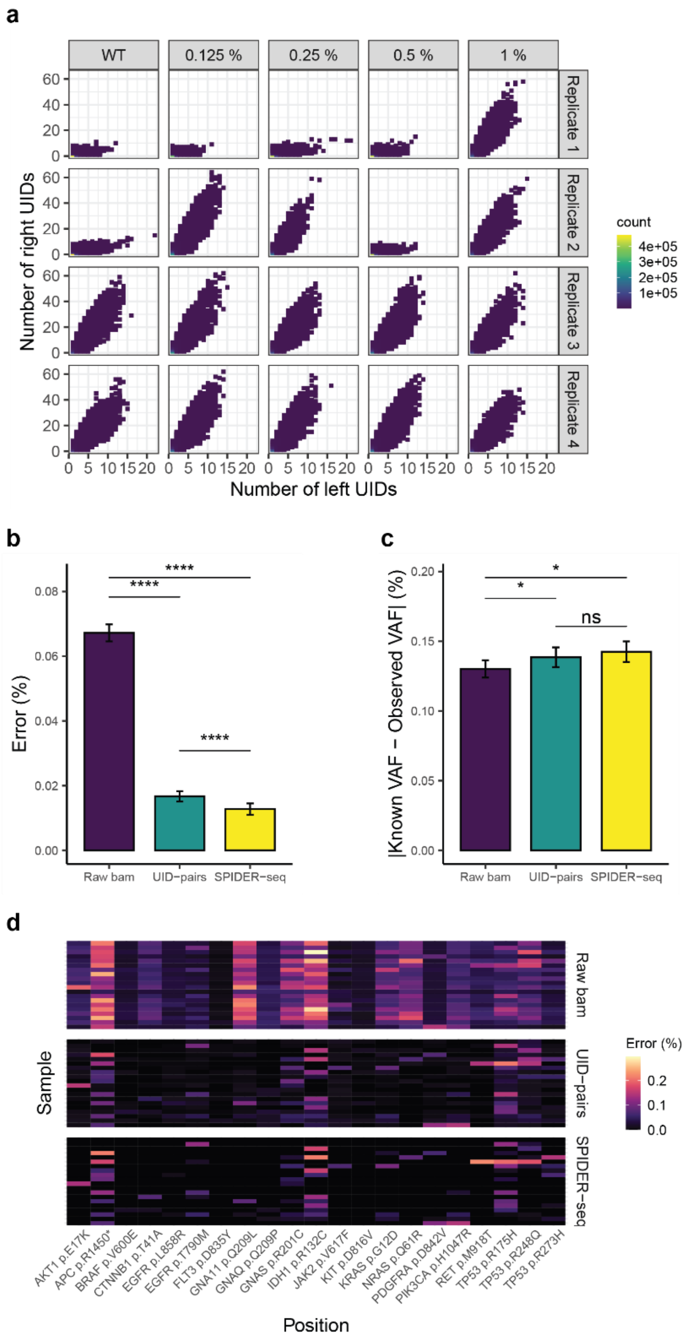


**Supplementary Figure 15.** Analysis of data from experiments using hybrid capture libraries. (**a**) Distribution of cluster size. (**b**) Comparison of error rate (%) across the following methods: base count from raw bam files, base count using UID-pairs, and base count using clusters (SPIDER-seq). Methods were compared using the Wilcoxon signed rank test ($P = 2.80 \times 10^{-67}$ between raw bam and SPIDER-seq method, $P = 4.13 \times 10^{-65}$ between raw bam and UID-pairs, and $P = 3.41 \times 10^{-21}$ between UID-pairs and SPIDER-seq method). (**c**) Absolute difference between known and observed VAF ($P = 4.02 \times 10^{-2}$ between raw bam and UID-pairs, $P = 1.44 \times 10^{-2}$ between raw bam and SPIDER-seq, and $P = 1.04 \times 10^{-1}$ between SPIDER-seq and UID-pairs. Error bars represent the standard error of the mean. (**d**) Error rate (%) across positions. Error rates were calculated by dividing the number of non-reference bases by the total number of bases at the targeted position. ns, not significant.

**Fig. S16**

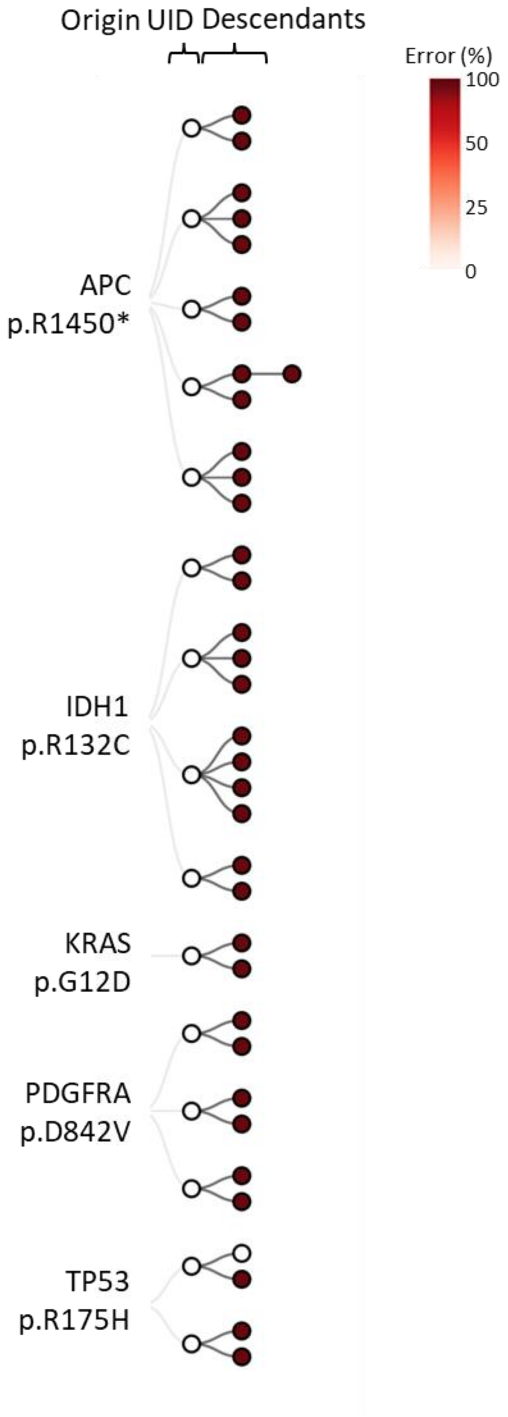


**Supplementary Figure 16.** Phylogenetic tree of clusters showing non-reference genotypes observed from analysis of a hybridization capture sample (WT, replicate 1).

**Fig. S17**

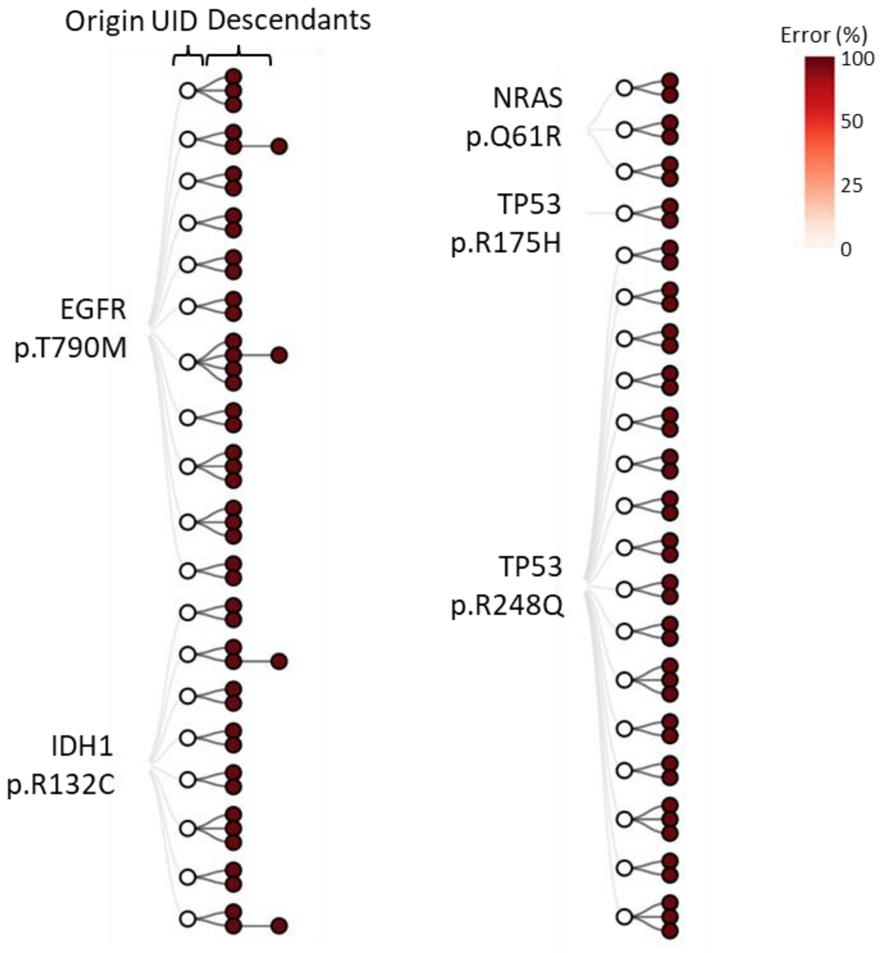


161
162
163 **Supplementary Figure 17.** Phylogenetic tree of clusters showing non-reference genotypes
164 observed from analysis of a hybridization capture sample (WT, replicate 2).
165

**Fig. S18**

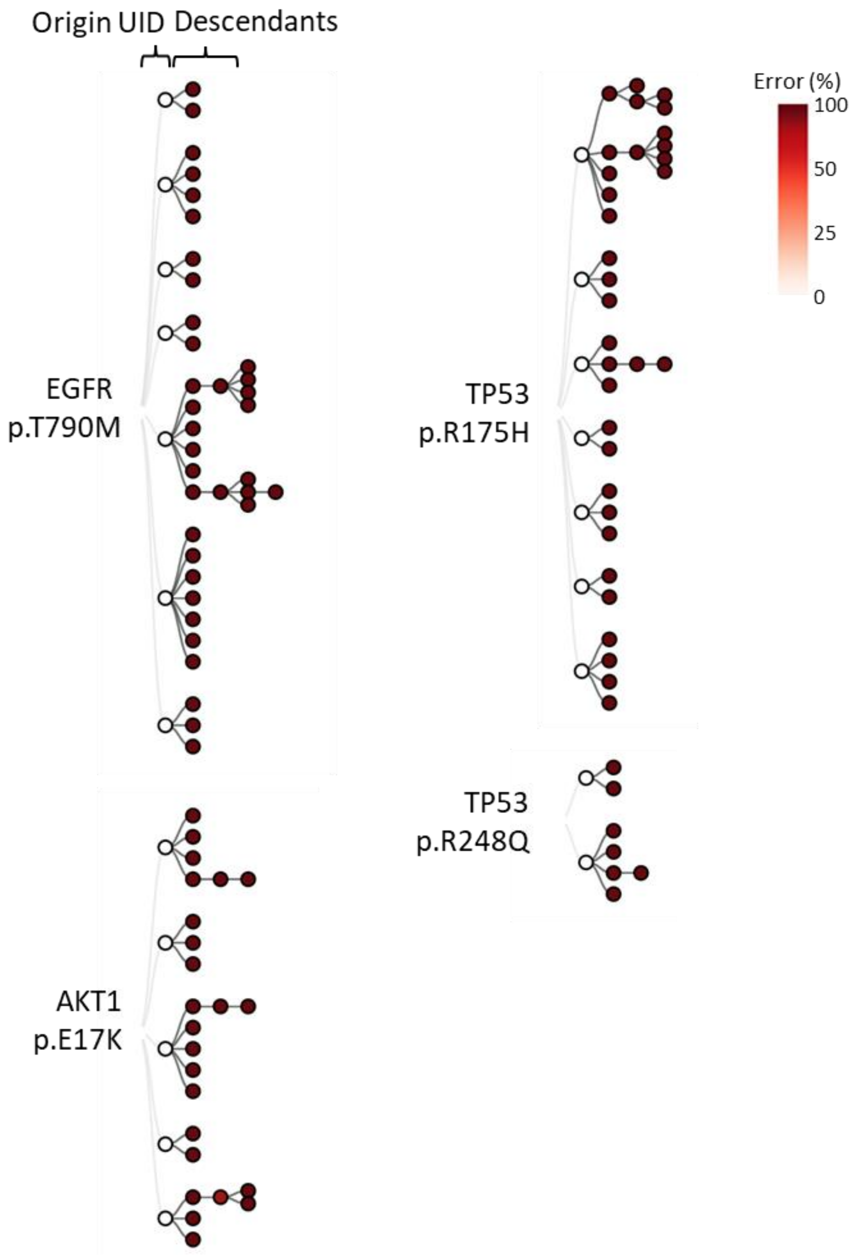


**Supplementary Figure 18.** Phylogenetic tree of clusters showing non-reference genotypes observed from analysis of a hybridization capture sample (WT, replicate 3).

**Supplementary Table 1.** Oligonucleotides used in this study. Yellow highlighting in sequences indicates degenerate bases (N = A, T, C, or G) and asterisks indicate phosphorothioate bonds.

| **Name** | **Sequence** |
|---|---|
| *For degenerate barcode content* | |
| *BRAF_N12* | actgttttcctttacttactacacctcagATATATTTCTTCATGAAGACCTCACAGTAAAAATAGGTGANNNNNNTCTAGCTACAGAGAAATCTCGATNNNNNNGGTCCCATCAGTTTGAACAGTTGTCTGGATCCATTTTGTGGATGgtaagaattgaggctatttttccac |
| *Primer for first round of amplification (UID-tagging amplification)* | |
| *NRAS_Q61_P5* | CACTCTTTCCCTACACGACGCTCTTCCGATCTTCGGTCACTTAGGANNNNANNNNGNNNNCNNNNATAGATGGTGAAACCTGTTTGTTGG |
| *KRAS_G12_P5* | CACTCTTTCCCTACACGACGCTCTTCCGATCTCGAGAGTTGGATGCTNNNNTNNNNANNNNGNNNNTATTATAAGGCCTGCTGAAAATG |
| *CTNNB1_T41_P5* | CACTCTTTCCCTACACGACGCTCTTCCGATCTGCATCAATGCCGTCANNNNCNNNNTNNNNANNNNCAACAGTCTTACCTGGACTCTGG |
| *JAK2_V617_P5* | CACTCTTTCCCTACACGACGCTCTTCCGATCTAGGTGGCGAACCTNNNNGNNNNCNNNNTNNNNAAGCTTTCTCACAAGCATTTGGTTT |
| *PDGFRA_D842_P5* | CACTCTTTCCCTACACGACGCTCTTCCGATCTTGCACTAACGATCCANNNNANNNNGNNNNCNNNNGCACAAGGAAAAATTGTGAAGAT |
| *PIK3CA_H1047_P5* | CACTCTTTCCCTACACGACGCTCTTCCGATCTCTCACTCCTCCAGTCNNNNCNNNNTNNNNANNNNAACTGAGCAAGAGGCTTTGG |
| *EGFR_T790_P5* | CACTCTTTCCCTACACGACGCTCTTCCGATCTCACTTACTCCGAACCNNNNANNNNGNNNNCNNNNGCAGGTACTGGGAGCCAAT |
| *EGFR_L858_P5* | CACTCTTTCCCTACACGACGCTCTTCCGATCTCAGAAGTGTGTGAGCNNNNANNNNGNNNNCNNNNGCAGCATGTCAAGATCACAGATT |
| *EGFR_ex19del_P5* | CACTCTTTCCCTACACGACGCTCTTCCGATCTCTTCAACTGATAGCGNNNNTNNNNANNNNGNNNNGAAAGTTAAAATTCCCGTCGCTAT |
| *BRAF_V600_P5* | CACTCTTTCCCTACACGACGCTCTTCCGATCTGACTTGTTCAGGATTNNNNTNNNNANNNNGNNNNTGAAGACCTCACAGTAAAAATAG |
| *NRAS_Q61_P7* | GACTGGAGTTCAGACGTGTGCTCTTCCGATCTAACGAGGTCTACTTCNNNNANNNNGNNNNCNNNNATGTATTGGTCTCTCATGGCA |
| *KRAS_G12_P7* | GACTGGAGTTCAGACGTGTGCTCTTCCGATCTGAACCGTACTCGTTCNNNNTNNNNANNNNGNNNNTATCGTCAAGGCACTCTT |
| *CTNNB1_T41_P7* | GACTGGAGTTCAGACGTGTGCTCTTCCGATCTGGCTTAAGGATCCAGNNNNCNNNNTNNNNANNNNCAGGATTGCCTTTACCACTCA |
| *JAK2_V617_P7* | GACTGGAGTTCAGACGTGTGCTCTTCCGATCTCCAGTCAGTGCTCNNNNGNNNNCNNNNTNNNNAGAAAGGCATTAGAAAGCCTGTAGTT |
| *PDGFRA_D842_P7* | GACTGGAGTTCAGACGTGTGCTCTTCCGATCTGAGAAGTTGCTCGAGNNNNANNNNGNNNNCNNNNAGGGAAGTGAGGACGTACACTG |

| | |
|---|---|
| *PIK3CA_H1047_P7* | GACTGGAGTTCAGACGTGTGCTCTTCCGATCTGCTTGTCTGAGTAGTNNNNCNNNNTNNNNANNNNCATTTTTGTTGTCCAGCCACC |
| *EGFR_T790_P7* | GACTGGAGTTCAGACGTGTGCTCTTCCGATCTCCATAGAGAACCAACNNNNTNNNNANNNNGNNNNGCATCTGCCTCACCTCCA |
| *EGFR_L858_P7* | GACTGGAGTTCAGACGTGTGCTCTTCCGATCTAGTGTATGGATACCNNNNANNNNGNNNNCNNNNCCTCCTTCTGCATGGTATTCTTTCT |
| *EGFR_ex19del_P7* | GACTGGAGTTCAGACGTGTGCTCTTCCGATCTTGCAAGTCGTAGACTNNNNTNNNNANNNNGNNNNAAAGCAGAAACTCACATCGA |
| *BRAF_V600_P7* | GACTGGAGTTCAGACGTGTGCTCTTCCGATCTTAGGTATCCTAAGCGNNNNTNNNNANNNNGNNNNATGGATCCAGACAACTGTTC |
| *Primers for amplification of hybridization capture library* | |
| NEBNext-i5-N5_1 | AATGATACGGCGACCACCGAGATCTACACGGCNNNNNACACTCTTTCCCTACACGACGCTCTTCCGATC*T |
| NEBNext-i5-N5_2 | AATGATACGGCGACCACCGAGATCTACACTCTNNNNNACACTCTTTCCCTACACGACGCTCTTCCGATC*T |
| NEBNext-i5-N5_3 | AATGATACGGCGACCACCGAGATCTACACCTANNNNNACACTCTTTCCCTACACGACGCTCTTCCGATC*T |
| NEBNext-i5-N5_4 | AATGATACGGCGACCACCGAGATCTACACAAGNNNNNACACTCTTTCCCTACACGACGCTCTTCCGATC*T |
| NEBNext-i7-N5_1 | CAAGCAGAAGACGGCATACGAGATTTGNNNNNGTGACTGGAGTTCAGACGTGTGCTCTTCCGATC*T |
| NEBNext-i7-N5_2 | CAAGCAGAAGACGGCATACGAGATGGTNNNNNGTGACTGGAGTTCAGACGTGTGCTCTTCCGATC*T |
| NEBNext-i7-N5_3 | CAAGCAGAAGACGGCATACGAGATCACNNNNNGTGACTGGAGTTCAGACGTGTGCTCTTCCGATC*T |
| NEBNext-i7-N5_4 | CAAGCAGAAGACGGCATACGAGATACANNNNNGTGACTGGAGTTCAGACGTGTGCTCTTCCGATC*T |

**Supplementary Table 2.** Materials used for this study.

| Product name | Material no. | Provider | Description |
|---|---|---|---|
| *cfDNA reference genomic DNA* | | | |
| Seraseq™ ctDNA Mutation Mix v2 WT | 0710-0144 | SeraCare Life Sciences | ctDNA model (Human, AF= 0%) |
| Seraseq™ ctDNA Mutation Mix v2 AF0.125% | 0710-0143 | SeraCare Life Sciences | ctDNA model (Human, AF = 0.125%) |
| Seraseq™ ctDNA Mutation Mix v2 AF0.25% | 0710-0142 | SeraCare Life Sciences | ctDNA model (Human, AF = 0.25%) |
| Seraseq™ ctDNA Mutation Mix v2 AF0.5% | 0710-0141 | SeraCare Life Sciences | ctDNA model (Human, AF = 0.5%) |
| Seraseq™ ctDNA Mutation Mix v2 AF1% | 0710-0140 | SeraCare Life Sciences | ctDNA model (Human, AF = 1%) |
| *Polymerases* | | | |
| HotStart PCR kit, with dNTPs | 07958897001 | Roche | KAPA polymerase |
| Phusion High-Fidelity DNA Polymerase | M0530S | NEB | Phusion polymerase |
| QIAGEN Multiplex PCR kit | 206143 | QIAGEN | Qiagen multiplex Taq polymerase |
| *Purification* | | | |
| AMPure XP | A63881 | BECKMAN COULTER | PCR cleanup kit for hybridization capture library |
| MinElute Gel Extraction kit | 28606 | QIAGEN | Purification kit of amplicon library |
| *Enzymes for preparation of hybridization capture library* | | | |
| 5× ER/A-Tailing Enzyme Mix | Y9420L | Enzymatics | Enzyme mix for end repair and A-tailing reaction |
| WGS Ligase | L6030-W-L | Enzymatics | Ligation of NGS adaptor |
| USER Enzyme | M5505S | NEB | Cleavage of uracil in the NEBNext adaptor |

**Supplementary Table 3.** Amount of cfDNA reference standard used in this study (hGE = haploid genome equivalent).

| Product name | Description | Concentration (ng/µl) | Experiments targeting *BRAF* | | Multiplex PCR experiment | | Hybrid capture experiments | |
|---|---|---|---|---|---|---|---|---|
| | | | ng | hGEs | ng | hGEs | ng | hGEs |
| Seraseq™ ctDNA Mutation Mix v2 WT | ctDNA model (Human, AF = 0%) | 15.6 | 15.6 | 4727 | 31.2 | 9455 | 31.2 | 9455 |
| Seraseq™ ctDNA Mutation Mix v2 AF0.125% | ctDNA model (Human, AF = 0.125%) | 15.8 | 15.8 | 4788 | 31.6 | 9576 | 31.6 | 9576 |
| Seraseq™ ctDNA Mutation Mix v2 AF0.25% | ctDNA model (Human, AF = 0.25%) | 13.9 | Not used | Not used | 27.8 | 8424 | 27.8 | 8424 |
| Seraseq™ ctDNA Mutation Mix v2 AF0.5% | ctDNA model (Human, AF = 0.5%) | 14.8 | 14.8 | 4485 | Not used | Not used | 29.6 | 8970 |
| Seraseq™ ctDNA Mutation Mix v2 AF1% | ctDNA model (Human, AF = 1%) | 12.2 | 12.2 | 3697 | Not used | Not used | 24.4 | 7394 |

**Supplementary Table 4.** The number of paired reads in the raw sequencing data for each sample.

| Experiment | Sample | Raw Reads |
|---|---|---:|
| oligonucleotide model | KP | 479,139,565 |
| | PH | 130,497,025 |
| | QM | 143,081,520 |
| Single locus | AF=0.125%-rep1 | 1,166,425 |
| | AF=0.125%-rep2 | 1,545,114 |
| | AF=0.125%-rep3 | 1,050,000 |
| | AF=0.5%-rep1 | 726,718 |
| | AF=0.5%-rep2 | 1,133,952 |
| | AF=0.5%-rep3 | 1,453,057 |
| | AF=1%-rep1 | 831,669 |
| | AF=1%-rep2 | 1,118,431 |
| | AF=1%-rep3 | 1,279,384 |
| | WT-rep1 | 614,570 |
| | WT-rep2 | 607,044 |
| | WT-rep3 | 477,585 |
| Multiple loci | AF=0.125%-rep1 | 229,589,906 |
| | AF=0.125%-rep2 | 220,966,519 |
| | AF=0.25%-rep1 | 212,625,203 |
| | AF=0.25%-rep2 | 229,063,887 |
| | WT-rep1 | 202,247,540 |
| | WT-rep2 | 209,957,751 |
| Hybridization | AF=0.125%-rep1 | 61,498,853 |
| | AF=0.125%-rep2 | 68,405,876 |
| | AF=0.125%-rep3 | 87,729,231 |
| | AF=0.125%-rep4 | 87,624,626 |
| | AF=0.25%-rep1 | 69,374,208 |
| | AF=0.25%-rep2 | 63,885,832 |
| | AF=0.25%-rep3 | 66,178,548 |
| | AF=0.25%-rep4 | 95,592,013 |
| | AF=0.5%-rep1 | 86,884,340 |
| | AF=0.5%-rep2 | 60,350,647 |
| | AF=0.5%-rep3 | 73,761,503 |
| | AF=0.5%-rep4 | 72,399,968 |
| | AF=1%-rep1 | 85,598,188 |
| | AF=1%-rep2 | 75,437,028 |
| | AF=1%-rep3 | 85,840,086 |
| | AF=1%-rep4 | 86,704,459 |
| | WT-rep1 | 69,103,422 |
| | WT-rep2 | 59,833,017 |
| | WT-rep3 | 77,115,432 |
| | WT-rep4 | 81,169,092 |

**Supplementary Table 5.** Number of reads, UID-pairs, CIDs, and barcode contents for each experiment.

| | KP | QM | PH |
|---|---|---|---|
| Trimmed paired-reads | 17,379,861 | 36,596,076 | 50,555,163 |
| UID-pairs | 1,280,164 | 2,249,912 | 2,205,754 |
| Used UID-pairs | 129,002 | 185,015 | 132,637 |
| Obtained CIDs | 58,114 | 93,525 | 65,438 |
| Number of contents | 2,888 | 3,526 | 2,560 |

**Supplementary Table 6.** Base distribution of the *BRAF* p.V600 locus. Each base was counted from raw data and consensus sequences based on CID and UIDs.

| Position (GRCh38) | Identifier | Variant allele frequency (%) | Replicate | A | T | C | G |
|---|---|---|---|---|---|---|---|
| chr7: 140,753,336 | CID | 0 | 1 | 21,025 | 0 | 0 | 0 |
| | | | 2 | 29,546 | 0 | 0 | 0 |
| | | | 3 | 14,854 | 4 | 0 | 0 |
| | | 0.125 | 1 | 73,245 | 42 | 0 | 0 |
| | | | 2 | 54,996 | 64 | 0 | 0 |
| | | | 3 | 58,247 | 40 | 0 | 0 |
| | | 0.5 | 1 | 66,458 | 357 | 0 | 0 |
| | | | 2 | 43,091 | 165 | 0 | 0 |
| | | | 3 | 40,267 | 190 | 0 | 0 |
| | | 1 | 1 | 26,576 | 193 | 0 | 0 |
| | | | 2 | 36,199 | 274 | 0 | 1 |
| | | | 3 | 47,240 | 585 | 0 | 10 |
| | UIDs | 0 | 1 | 104,381 | 6 | 0 | 15 |
| | | | 2 | 142,656 | 9 | 0 | 7 |
| | | | 3 | 79,283 | 27 | 1 | 3 |
| | | 0.125 | 1 | 390,596 | 202 | 1 | 36 |
| | | | 2 | 281,331 | 312 | 2 | 28 |
| | | | 3 | 331,816 | 294 | 2 | 34 |
| | | 0.5 | 1 | 328,937 | 1,765 | 1 | 37 |
| | | | 2 | 213,017 | 831 | 1 | 25 |
| | | | 3 | 194,835 | 960 | 0 | 15 |
| | | 1 | 1 | 150,492 | 1,186 | 5 | 12 |
| | | | 2 | 177,541 | 1,378 | 1 | 12 |
| | | | 3 | 252,082 | 3,214 | 1 | 58 |
| | No identifier (raw data) | 0 | 1 | 251,660 | 34 | 1 | 38 |
| | | | 2 | 372,229 | 64 | 8 | 28 |
| | | | 3 | 185,022 | 87 | 4 | 17 |
| | | 0.125 | 1 | 1169,451 | 680 | 10 | 151 |
| | | | 2 | 837,196 | 1,022 | 213 | 108 |
| | | | 3 | 734,586 | 713 | 5 | 107 |
| | | 0.5 | 1 | 795,916 | 4,335 | 66 | 101 |
| | | | 2 | 599,347 | 2,461 | 7 | 90 |
| | | | 3 | 518,203 | 2,600 | 2 | 72 |
| | | 1 | 1 | 319,169 | 2,556 | 13 | 40 |
| | | | 2 | 458,381 | 3,632 | 10 | 64 |
| | | | 3 | 798,129 | 10,239 | 11 | 241 |

**Supplementary Table 7.** Base distribution of the positions near the *BRAF* p.V600 locus from CID-based consensus.

| Position (GRCh38) | Variant allele frequency (%) | Replicate | A | T | C | G |
|---|---|---|---|---|---|---|
| chr7:140,753,332 | 0 | 1 | 0 | 21,031 | 0 | 0 |
| | | 2 | 0 | 29,552 | 0 | 0 |
| | | 3 | 0 | 14,857 | 0 | 0 |
| | 0.125 | 1 | 0 | 73,293 | 0 | 0 |
| | | 2 | 0 | 55,062 | 0 | 0 |
| | | 3 | 0 | 58,302 | 0 | 0 |
| | 0.5 | 1 | 0 | 66,825 | 0 | 0 |
| | | 2 | 0 | 43,260 | 0 | 0 |
| | | 3 | 0 | 40,459 | 0 | 0 |
| | 1 | 1 | 0 | 26,777 | 0 | 0 |
| | | 2 | 0 | 36,484 | 0 | 0 |
| | | 3 | 0 | 47,845 | 0 | 0 |
| chr7:140,753,333 | 0 | 1 | 0 | 21,030 | 0 | 0 |
| | | 2 | 0 | 29,548 | 0 | 0 |
| | | 3 | 0 | 14,853 | 0 | 0 |
| | 0.125 | 1 | 0 | 73,284 | 0 | 0 |
| | | 2 | 0 | 55,063 | 0 | 0 |
| | | 3 | 0 | 58,283 | 0 | 0 |
| | 0.5 | 1 | 0 | 66,809 | 0 | 0 |
| | | 2 | 0 | 43,253 | 0 | 0 |
| | | 3 | 0 | 40,449 | 0 | 0 |
| | 1 | 1 | 0 | 26,774 | 0 | 0 |
| | | 2 | 0 | 36,477 | 0 | 0 |
| | | 3 | 0 | 47,827 | 0 | 0 |
| chr7:140,753,334 | 0 | 1 | 0 | 21,020 | 4 | 0 |
| | | 2 | 0 | 29,551 | 0 | 0 |
| | | 3 | 0 | 14,858 | 0 | 0 |
| | 0.125 | 1 | 0 | 73,294 | 0 | 0 |
| | | 2 | 0 | 55,058 | 0 | 0 |
| | | 3 | 0 | 58,289 | 0 | 0 |
| | 0.5 | 1 | 0 | 66,825 | 0 | 0 |
| | | 2 | 0 | 43,265 | 0 | 0 |
| | | 3 | 0 | 40,460 | 0 | 0 |
| | 1 | 1 | 0 | 26,776 | 0 | 0 |
| | | 2 | 0 | 36,471 | 0 | 0 |
| | | 3 | 0 | 47,837 | 0 | 0 |
| chr7:140,753,335 | 0 | 1 | 0 | 0 | 21,027 | 0 |
| | | 2 | 0 | 0 | 29,550 | 0 |
| | | 3 | 6 | 0 | 14,853 | 0 |

| | | | | | | |
|---|---|---|---|---|---|---|
| | 0.125 | 1 | 3 | 0 | 73,292 | 0 |
| | | 2 | 38 | 0 | 55,029 | 0 |
| | | 3 | 9 | 6 | 58,282 | 0 |
| | 0.5 | 1 | 14 | 1 | 66,814 | 0 |
| | | 2 | 0 | 0 | 43,265 | 0 |
| | | 3 | 0 | 0 | 40,460 | 0 |
| | 1 | 1 | 0 | 0 | 26,769 | 0 |
| | | 2 | 30 | 0 | 36,439 | 0 |
| | | 3 | 0 | 1 | 47,842 | 0 |
| chr7:140,753,337 | 0 | 1 | 10 | 0 | 21,016 | 0 |
| | | 2 | 0 | 0 | 29,551 | 0 |
| | | 3 | 0 | 0 | 14,853 | 0 |
| | 0.125 | 1 | 0 | 2 | 73,290 | 1 |
| | | 2 | 0 | 0 | 55,060 | 0 |
| | | 3 | 0 | 0 | 58,295 | 0 |
| | 0.5 | 1 | 0 | 4 | 66,816 | 0 |
| | | 2 | 15 | 0 | 43,255 | 0 |
| | | 3 | 0 | 12 | 40,448 | 0 |
| | 1 | 1 | 0 | 0 | 26,780 | 0 |
| | | 2 | 0 | 0 | 36,478 | 0 |
| | | 3 | 0 | 0 | 47,837 | 0 |
| chr7:140,753,338 | 0 | 1 | 0 | 21,023 | 1 | 0 |
| | | 2 | 0 | 29,548 | 0 | 0 |
| | | 3 | 0 | 14,860 | 0 | 0 |
| | 0.125 | 1 | 0 | 73,285 | 0 | 0 |
| | | 2 | 0 | 55,050 | 0 | 0 |
| | | 3 | 0 | 58,286 | 0 | 0 |
| | 0.5 | 1 | 0 | 66,805 | 10 | 0 |
| | | 2 | 0 | 43,264 | 1 | 0 |
| | | 3 | 0 | 40,459 | 0 | 0 |
| | 1 | 1 | 0 | 26,773 | 0 | 0 |
| | | 2 | 0 | 36,477 | 0 | 0 |
| | | 3 | 0 | 47,842 | 0 | 0 |
| chr7:140,753,339 | 0 | 1 | 0 | 0 | 0 | 21,028 |
| | | 2 | 24 | 0 | 0 | 29,522 |
| | | 3 | 0 | 0 | 0 | 14,861 |
| | 0.125 | 1 | 8 | 0 | 0 | 73,282 |
| | | 2 | 0 | 0 | 0 | 55,061 |
| | | 3 | 0 | 0 | 0 | 58,279 |
| | 0.5 | 1 | 0 | 11 | 0 | 66,808 |
| | | 2 | 2 | 17 | 0 | 43,245 |
| | | 3 | 0 | 0 | 0 | 40,461 |
| | 1 | 1 | 0 | 11 | 0 | 26,763 |
| | | 2 | 1 | 0 | 0 | 36,475 |
| | | 3 | 0 | 21 | 0 | 47,821 |

| | | | | | | |
|---|---|---|---|---|---|---|
| chr7:140,753,340 | 0 | 1 | 0 | 21,029 | 0 | 0 |
| | | 2 | 0 | 29,551 | 0 | 0 |
| | | 3 | 0 | 14,860 | 0 | 0 |
| | 0.125 | 1 | 0 | 73,289 | 4 | 0 |
| | | 2 | 0 | 55,063 | 0 | 0 |
| | | 3 | 0 | 58,296 | 0 | 0 |
| | 0.5 | 1 | 0 | 66,828 | 0 | 0 |
| | | 2 | 0 | 43,265 | 0 | 0 |
| | | 3 | 0 | 40,454 | 0 | 0 |
| | 1 | 1 | 0 | 26,772 | 0 | 0 |
| | | 2 | 0 | 36,475 | 0 | 0 |
| | | 3 | 0 | 47,843 | 0 | 0 |

203

**Supplementary Table 8.** Target list for multiplex PCR experiment.

| Target | Mutation type | HGVS_ nomenclature | Mutation position (GRCh38) | Strand | Amplicon size |
|---|---|---|---|---|---|
| *NRAS* (p.Q61R) | Substitution | c.182A>G | chr1:114,713,908 | − | 78 |
| *KRAS* (p.G12D) | Substitution | c.35G>A | chr12:25,245,350 | − | 81 |
| *CTNNB1* (p.T41A) | Substitution | c.121A>G | chr3:41,224,633 | + | 77 |
| *JAK2* (p.V617F) | Substitution | c.1849G>T | chr9:5,073,770 | + | 90 |
| *PDGFRA* (p.D842V) | Substitution | c.2525A>T | chr4:54,285,926 | + | 100 |
| *PIK3CA* (p.H1047R) | Substitution | c.3140A>G | chr3:179,234,297 | + | 74 |
| *EGFR* (p.T790M) | Substitution | c.2369C>T | chr7:55,181,378 | + | 106 |
| *EGFR* (p.L858R) | Substitution | c.2573T>G | chr7:55,191,822 | + | 76 |
| *BRAF* (p.V600E) | Substitution | c.1799T>A | chr7:140,753,336 | − | 94 |
| *EGFR* (p.E746_A750 del ELREA) | Deletion | c.2236 _2250del15 | chr7:55,174,773 -55,174,787 | + | 89 |

**Supplementary Table 9.** Target list for hybridization capture.

| Target | Mutation type | HGVS_nomenclature | Mutation position (GRCh38) | Strand |
|---|---|---|---|---|
| NRAS p.Q61R | Substitution | c.182A>G | chr1:114,713,908 | − |
| RET p.M918T | Substitution | c.2753T>C | chr10:43,121,968 | + |
| ATM p.C353fs*5 | Deletion | c.1058_1059delGT | chr11:108,247,120-108,247,121 | + |
| KRAS p.G12D | Substitution | c.35G>A | chr12:25,245,350 | − |
| FLT3 p.D835Y | Substitution | c.2503G>T | chr13:28,018,505 | − |
| AKT1 p.E17K | Substitution | c.49G>A | chr14:104,780,214 | − |
| ERBB2 p.A775_G776insYVMA | Insertion | c.2324_2325ins12 | chr17:39,724,742-39,724,743 | + |
| TP53 p.R175H | Substitution | c.524G>A | chr17:7,675,088 | − |
| TP53 p.R248Q | Substitution | c.743G>A | chr17:7,674,220 | − |
| TP53 p.R273H | Substitution | c.818G>A | chr17:7,673,802 | − |
| GNA11 p.Q209L | Substitution | c.626A>T | chr19:3,118,944 | + |
| IDH1 p.R132C | Substitution | c.394C>T | chr2:208,248,389 | − |
| GNAS p.R201C | Substitution | c.601C>T | chr20:58,909,365 | + |
| CTNNB1 p.T41A | Substitution | c.121A>G | chr3:41,224,633 | + |
| FOXL2 p.C134W | Substitution | c.402C>G | chr3:138,946,321 | − |
| PIK3CA p.E545K | Substitution | c.1633G>A | chr3:179,218,303 | + |
| PIK3CA p.H1047R | Substitution | c.3140A>G | chr3:179,234,297 | + |
| FGFR3 p.S249C | Substitution | c.746C>G | chr4:1,801,841 | + |
| KIT p.D816V | Substitution | c.2447A>T | chr4:54,733,155 | + |
| PDGFRA p.D842V | Substitution | c.2525A>T | chr4:54,285,926 | + |
| APC p.R1450* | Substitution | c.4348C>T | chr5:112,839,942 | + |
| EGFR p.E746_A750delELREA | Deletion | c.2236_2250del15 | chr7:55,174,773-55,174,787 | + |
| EGFR p.D770_N771insG | Insertion | c.2310_2311insGGT | chr7:55,181,319-55,181,320 | + |
| EGFR p.L858R | Substitution | c.2573T>G | chr7:55,191,822 | + |
| BRAF p.V600E | Substitution | c.1799T>A | chr7:140,753,336 | − |
| EGFR p.T790M | Substitution | c.2369C>T | chr7:55,181,378 | + |
| GNAQ p.Q209P | Substitution | c.626A>C | chr9:77,794,572 | − |
| JAK2 p.V617F | Substitution | c.1849G>T | chr9:5,073,770 | + |

**Supplementary Table 10.** Coverage per experiment.

| Replicate | Variant Allele Frequency (%) | replicate 1 | replicate 2 | replicate 3 | replicate 4 |
|---|---|---|---|---|---|
| AKT1-p.E17K | 0 | 385087 | 290282 | 435919 | 411243 |
| APC-p.R1450* | | 271004 | 204143 | 323543 | 326981 |
| ATM-p.C353fs*5 | | 266194 | 196108 | 274922 | 280229 |
| BRAF-p.V600E | | 326257 | 232642 | 310381 | 322605 |
| CTNNB1-p.T41A | | 577006 | 432372 | 605078 | 612902 |
| EGFR-p.D770_N771insG | | 670323 | 548045 | 653662 | 688472 |
| EGFR-p.E746_A750delELREA | | 235825 | 180339 | 260573 | 258897 |
| EGFR-p.L858R | | 752832 | 563805 | 690438 | 777531 |
| EGFR-p.T790M | | 742562 | 615392 | 739939 | 770818 |
| ERBB2-p.A775_G776insYVMA | | 715691 | 580868 | 832360 | 902435 |
| FGFR3-p.S249C | | 51687 | 40553 | 46463 | 51604 |
| FLT3-p.D835Y | | 434036 | 323959 | 415116 | 418313 |
| FOXL2-p.C134W | | 88443 | 78974 | 73363 | 80827 |
| GNA11-p.Q209L | | 550798 | 453012 | 648473 | 639805 |
| GNAQ-p.Q209P | | 324003 | 270423 | 309730 | 335105 |
| GNAS-p.R201C | | 273720 | 216435 | 293799 | 325356 |
| IDH1-p.R132C | | 369479 | 276122 | 361381 | 376629 |
| JAK2-p.V617F | | 402254 | 303246 | 370567 | 371570 |
| KIT-p.D816V | | 417346 | 330100 | 414802 | 448430 |
| KRAS-p.G12D | | 493407 | 349848 | 418577 | 466475 |
| NRAS-p.Q61R | | 306714 | 219640 | 267041 | 282955 |
| PDGFRA-p.D842V | | 500706 | 368601 | 531649 | 517931 |
| PIK3CA-p.E545K | | 44778 | 35115 | 38926 | 44164 |
| PIK3CA-p.H1047R | | 433206 | 327090 | 434958 | 478961 |
| RET-p.M918T | | 346406 | 279298 | 338412 | 335418 |
| TP53-p.R175H | | 834909 | 607283 | 751572 | 822903 |
| TP53-p.R248Q | | 763062 | 601957 | 826733 | 811083 |
| TP53-p.R273H | | 497425 | 390444 | 495590 | 509962 |
| AKT1-p.E17K | 0.125 | 291818 | 358964 | 458123 | 353622 |
| APC-p.R1450* | | 177596 | 230609 | 276249 | 210585 |
| ATM-p.C353fs*5 | | 148836 | 132578 | 179457 | 137295 |
| BRAF-p.V600E | | 200284 | 155054 | 184913 | 169534 |
| CTNNB1-p.T41A | | 410072 | 421662 | 538555 | 437159 |
| EGFR-p.D770_N771insG | | 517063 | 598588 | 792973 | 611236 |
| EGFR-p.E746_A750delELREA | | 156402 | 191311 | 240321 | 182118 |
| EGFR-p.L858R | | 474021 | 542134 | 647430 | 507515 |

| | | | | | |
|---|---|---|---|---|---|
| EGFR-p.T790M | | 595735 | 643170 | 855499 | 641082 |
| ERBB2-p.A775_G776insYVMA | | 541722 | 649415 | 850544 | 680777 |
| FGFR3-p.S249C | | 43897 | 48860 | 62164 | 48843 |
| FLT3-p.D835Y | | 297980 | 310725 | 376299 | 308177 |
| FOXL2-p.C134W | | 63544 | 70176 | 99121 | 73442 |
| GNA11-p.Q209L | | 418689 | 497786 | 622246 | 561045 |
| GNAQ-p.Q209P | | 198962 | 176543 | 213609 | 173550 |
| GNAS-p.R201C | | 207709 | 223026 | 280587 | 226198 |
| IDH1-p.R132C | | 266667 | 240992 | 285963 | 245869 |
| JAK2-p.V617F | | 237045 | 197116 | 238961 | 203728 |
| KIT-p.D816V | | 258938 | 221485 | 278536 | 226706 |
| KRAS-p.G12D | | 295642 | 258166 | 316254 | 263426 |
| NRAS-p.Q61R | | 220865 | 207561 | 231387 | 209334 |
| PDGFRA-p.D842V | | 323232 | 380752 | 477192 | 375221 |
| PIK3CA-p.E545K | | 19987 | 18325 | 20301 | 18068 |
| PIK3CA-p.H1047R | | 279231 | 265601 | 323312 | 269603 |
| RET-p.M918T | | 223554 | 254192 | 304633 | 243818 |
| TP53-p.R175H | | 600662 | 680827 | 880584 | 666725 |
| TP53-p.R248Q | | 606878 | 715176 | 832819 | 708169 |
| TP53-p.R273H | | 348103 | 365668 | 455495 | 338875 |
| AKT1-p.E17K | 0.25 | 392849 | 110609 | 243588 | 409311 |
| APC-p.R1450* | | 297012 | 82005 | 240738 | 331858 |
| ATM-p.C353fs*5 | | 258058 | 74308 | 215021 | 315330 |
| BRAF-p.V600E | | 282463 | 83040 | 236819 | 343286 |
| CTNNB1-p.T41A | | 556474 | 153841 | 432184 | 598725 |
| EGFR-p.D770_N771insG | | 631933 | 184620 | 430576 | 700095 |
| EGFR-p.E746_A750delELREA | | 260333 | 82380 | 210421 | 312823 |
| EGFR-p.L858R | | 703631 | 194464 | 483343 | 758842 |
| EGFR-p.T790M | | 674471 | 196891 | 469134 | 730536 |
| ERBB2-p.A775_G776insYVMA | | 704764 | 203187 | 498778 | 756048 |
| FGFR3-p.S249C | | 55940 | 17963 | 30196 | 62708 |
| FLT3-p.D835Y | | 366447 | 103213 | 292675 | 425740 |
| FOXL2-p.C134W | | 98497 | 24573 | 55586 | 87501 |
| GNA11-p.Q209L | | 654086 | 176323 | 411163 | 686187 |
| GNAQ-p.Q209P | | 246198 | 76766 | 234460 | 332367 |
| GNAS-p.R201C | | 305811 | 82901 | 225473 | 346336 |
| IDH1-p.R132C | | 356785 | 106840 | 305187 | 420183 |
| JAK2-p.V617F | | 351406 | 101303 | 295524 | 441442 |
| KIT-p.D816V | | 377283 | 107499 | 322499 | 450291 |
| KRAS-p.G12D | | 375774 | 101249 | 316712 | 414388 |
| NRAS-p.Q61R | | 245353 | 68050 | 200976 | 271175 |
| PDGFRA-p.D842V | | 507389 | 135498 | 372308 | 560502 |
| PIK3CA-p.E545K | | 41348 | 12061 | 37015 | 50746 |

| | | | | | |
|---|---|---|---|---|---|
| PIK3CA-p.H1047R | | 368311 | 108111 | 332804 | 473388 |
| RET-p.M918T | | 297379 | 92500 | 244429 | 376182 |
| TP53-p.R175H | | 719675 | 196514 | 478048 | 795687 |
| TP53-p.R248Q | | 726627 | 209101 | 515337 | 794057 |
| TP53-p.R273H | | 460993 | 128856 | 342250 | 527136 |
| AKT1-p.E17K | 0.5 | 464440 | 219039 | 399452 | 477427 |
| APC-p.R1450* | | 335243 | 130947 | 258774 | 283888 |
| ATM-p.C353fs*5 | | 235149 | 113863 | 202403 | 230748 |
| BRAF-p.V600E | | 287184 | 138486 | 250961 | 282152 |
| CTNNB1-p.T41A | | 657466 | 285125 | 540398 | 589719 |
| EGFR-p.D770_N771insG | | 815756 | 373080 | 657086 | 750294 |
| EGFR-p.E746_A750delELREA | | 275097 | 117067 | 253407 | 272599 |
| EGFR-p.L858R | | 726918 | 396019 | 694977 | 755262 |
| EGFR-p.T790M | | 888161 | 418082 | 710061 | 820762 |
| ERBB2-p.A775_G776insYVMA | | 821922 | 418613 | 758721 | 828054 |
| FGFR3-p.S249C | | 60397 | 28499 | 60684 | 65622 |
| FLT3-p.D835Y | | 464201 | 220534 | 391916 | 426451 |
| FOXL2-p.C134W | | 112889 | 52105 | 82014 | 93911 |
| GNA11-p.Q209L | | 710920 | 353351 | 622261 | 660159 |
| GNAQ-p.Q209P | | 291534 | 135744 | 258966 | 281788 |
| GNAS-p.R201C | | 320095 | 156726 | 272577 | 311993 |
| IDH1-p.R132C | | 363335 | 194396 | 352872 | 385544 |
| JAK2-p.V617F | | 355087 | 168133 | 296777 | 332601 |
| KIT-p.D816V | | 391680 | 191215 | 324847 | 368422 |
| KRAS-p.G12D | | 418328 | 209253 | 363548 | 397659 |
| NRAS-p.Q61R | | 278688 | 148294 | 251401 | 275948 |
| PDGFRA-p.D842V | | 554472 | 247443 | 479176 | 538187 |
| PIK3CA-p.E545K | | 34384 | 18428 | 27464 | 32864 |
| PIK3CA-p.H1047R | | 392546 | 201238 | 335435 | 407163 |
| RET-p.M918T | | 343749 | 170832 | 303805 | 355676 |
| TP53-p.R175H | | 918946 | 447491 | 785769 | 899555 |
| TP53-p.R248Q | | 861374 | 440565 | 797500 | 903357 |
| TP53-p.R273H | | 526072 | 250382 | 464640 | 538217 |
| AKT1-p.E17K | 1 | 188185 | 264210 | 365880 | 346579 |
| APC-p.R1450* | | 161316 | 255094 | 289174 | 277891 |
| ATM-p.C353fs*5 | | 130400 | 185553 | 243775 | 254193 |
| BRAF-p.V600E | | 154927 | 222349 | 279540 | 268400 |
| CTNNB1-p.T41A | | 316912 | 440898 | 547876 | 563574 |
| EGFR-p.D770_N771insG | | 331354 | 499108 | 616120 | 596998 |
| EGFR-p.E746_A750delELREA | | 152286 | 218903 | 264943 | 233773 |
| EGFR-p.L858R | | 352950 | 547447 | 644850 | 606492 |
| EGFR-p.T790M | | 355540 | 534454 | 661214 | 637232 |
| ERBB2-p.A775_G776insYVMA | | 348986 | 540811 | 663555 | 631434 |

| | | | | | |
|---|---|---|---|---|---|
| FGFR3-p.S249C | | 21882 | 28292 | 43569 | 36833 |
| FLT3-p.D835Y | | 205494 | 310281 | 395321 | 356008 |
| FOXL2-p.C134W | | 41022 | 57483 | 85841 | 74818 |
| GNA11-p.Q209L | | 283654 | 368656 | 502975 | 490103 |
| GNAQ-p.Q209P | | 158219 | 217845 | 292753 | 262694 |
| GNAS-p.R201C | | 161962 | 227938 | 305396 | 271843 |
| IDH1-p.R132C | | 214241 | 314317 | 379620 | 367287 |
| JAK2-p.V617F | | 183674 | 265174 | 328553 | 334642 |
| KIT-p.D816V | | 211608 | 313664 | 380049 | 362399 |
| KRAS-p.G12D | | 217651 | 307948 | 406711 | 386587 |
| NRAS-p.Q61R | | 165336 | 213936 | 263044 | 261314 |
| PDGFRA-p.D842V | | 272141 | 384092 | 474566 | 484680 |
| PIK3CA-p.E545K | | 18982 | 26832 | 33639 | 34570 |
| PIK3CA-p.H1047R | | 234991 | 345097 | 437220 | 407976 |
| RET-p.M918T | | 188629 | 269911 | 328189 | 310859 |
| TP53-p.R175H | | 388012 | 531587 | 666912 | 606570 |
| TP53-p.R248Q | | 414160 | 532470 | 676403 | 668171 |
| TP53-p.R273H | | 252855 | 376991 | 453221 | 407341 |

**Supplementary Table 11.** Number of consensus reads for each experiment.

| Replicate | Variant Allele Frequency (%) | replicate 1 | replicate 2 | replicate 3 | replicate 4 |
|---|---|---|---|---|---|
| AKT1-p.E17K | 0 | 3712 | 6069 | 4504 | 2402 |
| APC-p.R1450* | | 2371 | 3477 | 4480 | 2778 |
| ATM-p.C353fs*5 | | 2246 | 3241 | 4102 | 2600 |
| BRAF-p.V600E | | 2675 | 3996 | 4096 | 2623 |
| CTNNB1-p.T41A | | 5971 | 8663 | 6252 | 3995 |
| EGFR-p.D770_N771insG | | 10694 | 18057 | 9728 | 5377 |
| EGFR-p.E746_A750delELREA | | 1615 | 2928 | 3212 | 1936 |
| EGFR-p.L858R | | 8444 | 11915 | 5062 | 3152 |
| EGFR-p.T790M | | 6902 | 11925 | 5671 | 3456 |
| ERBB2-p.A775_G776insYVMA | | 6436 | 9987 | 6549 | 3877 |
| FLT3-p.D835Y | | 3820 | 5547 | 5191 | 3124 |
| GNA11-p.Q209L | | 5485 | 8855 | 5717 | 3518 |
| GNAQ-p.Q209P | | 2717 | 4412 | 4564 | 3094 |
| GNAS-p.R201C | | 2292 | 4328 | 4263 | 2863 |
| IDH1-p.R132C | | 3271 | 4818 | 4752 | 3006 |
| JAK2-p.V617F | | 3678 | 5288 | 4749 | 2991 |
| KIT-p.D816V | | 3790 | 6331 | 5005 | 3369 |
| KRAS-p.G12D | | 4391 | 6321 | 5386 | 3723 |
| NRAS-p.Q61R | | 2986 | 4491 | 3606 | 2302 |
| PDGFRA-p.D842V | | 4574 | 6665 | 5005 | 3054 |
| PIK3CA-p.H1047R | | 3414 | 5649 | 5429 | 3678 |
| RET-p.M918T | | 2959 | 5131 | 4067 | 2366 |
| TP53-p.R175H | | 7598 | 11545 | 5733 | 3412 |
| TP53-p.R248Q | | 8372 | 12089 | 6732 | 3951 |
| TP53-p.R273H | | 5489 | 8573 | 4562 | 2729 |
| AKT1-p.E17K | 0.125 | 2754 | 12390 | 13531 | 9725 |
| APC-p.R1450* | | 1320 | 9631 | 10122 | 7687 |
| ATM-p.C353fs*5 | | 1239 | 6136 | 6719 | 5100 |
| BRAF-p.V600E | | 1548 | 6666 | 6733 | 5901 |
| CTNNB1-p.T41A | | 3977 | 16247 | 17678 | 12908 |
| EGFR-p.D770_N771insG | | 8022 | 28439 | 30457 | 22404 |
| EGFR-p.E746_A750delELREA | | 982 | 7370 | 7928 | 5850 |
| EGFR-p.L858R | | 5091 | 14222 | 14472 | 10879 |
| EGFR-p.T790M | | 5335 | 16845 | 18123 | 13157 |

| | | | | | |
|---|---|---|---|---|---|
| ERBB2-p.A775_G776insYVMA | | 4771 | 17509 | 18621 | 13692 |
| FLT3-p.D835Y | | 2487 | 11985 | 12541 | 9773 |
| GNA11-p.Q209L | | 4042 | 14758 | 15963 | 12635 |
| GNAQ-p.Q209P | | 1608 | 8102 | 8219 | 6391 |
| GNAS-p.R201C | | 1581 | 9896 | 10846 | 8397 |
| IDH1-p.R132C | | 2289 | 9829 | 10070 | 8126 |
| JAK2-p.V617F | | 2118 | 8388 | 8672 | 6879 |
| KIT-p.D816V | | 2281 | 9256 | 9780 | 7482 |
| KRAS-p.G12D | | 2650 | 11358 | 11706 | 8775 |
| NRAS-p.Q61R | | 1987 | 9185 | 8895 | 7380 |
| PDGFRA-p.D842V | | 2729 | 12704 | 13194 | 9788 |
| PIK3CA-p.H1047R | | 2106 | 10610 | 10952 | 8795 |
| RET-p.M918T | | 1728 | 10033 | 10598 | 7873 |
| TP53-p.R175H | | 5066 | 17591 | 18706 | 13839 |
| TP53-p.R248Q | | 6375 | 20340 | 21520 | 16063 |
| TP53-p.R273H | | 3591 | 12006 | 12911 | 9367 |
| AKT1-p.E17K | 0.25 | 5601 | 5788 | 3611 | 5923 |
| APC-p.R1450* | | 3569 | 4716 | 4485 | 6463 |
| ATM-p.C353fs*5 | | 2972 | 4517 | 4208 | 6561 |
| BRAF-p.V600E | | 3302 | 4623 | 4260 | 6441 |
| CTNNB1-p.T41A | | 7863 | 8690 | 6589 | 10049 |
| EGFR-p.D770_N771insG | | 13955 | 14115 | 9438 | 15281 |
| EGFR-p.E746_A750delELREA | | 2780 | 4368 | 3604 | 5471 |
| EGFR-p.L858R | | 9779 | 8683 | 5127 | 7913 |
| EGFR-p.T790M | | 8975 | 8523 | 5432 | 8524 |
| ERBB2-p.A775_G776insYVMA | | 8949 | 9577 | 5731 | 8994 |
| FLT3-p.D835Y | | 4341 | 5811 | 5084 | 7530 |
| GNA11-p.Q209L | | 8920 | 8525 | 5578 | 9115 |
| GNAQ-p.Q209P | | 2759 | 4666 | 4652 | 6843 |
| GNAS-p.R201C | | 3876 | 4989 | 4477 | 7110 |
| IDH1-p.R132C | | 4385 | 6166 | 5480 | 7819 |
| JAK2-p.V617F | | 4266 | 6126 | 5234 | 8294 |
| KIT-p.D816V | | 4808 | 6130 | 5560 | 7984 |
| KRAS-p.G12D | | 4621 | 6227 | 5873 | 7977 |
| NRAS-p.Q61R | | 3348 | 4289 | 3809 | 5317 |
| PDGFRA-p.D842V | | 6392 | 6923 | 5284 | 8222 |
| PIK3CA-p.H1047R | | 4205 | 6022 | 5588 | 8499 |
| RET-p.M918T | | 3672 | 5245 | 4152 | 6612 |

| | | | | | |
|---|---|---|---|---|---|
| TP53-p.R175H | | 9549 | 8340 | 5433 | 8634 |
| TP53-p.R248Q | | 10456 | 9925 | 6247 | 10446 |
| TP53-p.R273H | | 6785 | 6609 | 4915 | 7290 |
| AKT1-p.E17K | 0.5 | 4688 | 5114 | 7776 | 9929 |
| APC-p.R1450* | | 3130 | 2211 | 6700 | 8074 |
| ATM-p.C353fs*5 | | 2058 | 2048 | 5376 | 7019 |
| BRAF-p.V600E | | 2425 | 2541 | 5959 | 7521 |
| CTNNB1-p.T41A | | 6898 | 6577 | 11785 | 14003 |
| EGFR-p.D770_N771insG | | 13586 | 14158 | 19283 | 23017 |
| EGFR-p.E746_A750delELREA | | 1980 | 1919 | 5877 | 7074 |
| EGFR-p.L858R | | 7729 | 9411 | 10270 | 11468 |
| EGFR-p.T790M | | 8456 | 9299 | 11213 | 13213 |
| ERBB2-p.A775_G776insYVMA | | 7624 | 8527 | 11859 | 13735 |
| FLT3-p.D835Y | | 4024 | 4127 | 8802 | 10831 |
| GNA11-p.Q209L | | 6808 | 7754 | 10163 | 11756 |
| GNAQ-p.Q209P | | 2485 | 2273 | 6819 | 8422 |
| GNAS-p.R201C | | 2752 | 3189 | 6931 | 9197 |
| IDH1-p.R132C | | 3183 | 3696 | 8401 | 10215 |
| JAK2-p.V617F | | 3269 | 3011 | 7018 | 8485 |
| KIT-p.D816V | | 3658 | 3921 | 7295 | 9090 |
| KRAS-p.G12D | | 3855 | 4014 | 8950 | 10771 |
| NRAS-p.Q61R | | 2734 | 3069 | 6321 | 7699 |
| PDGFRA-p.D842V | | 5214 | 4955 | 9293 | 11393 |
| PIK3CA-p.H1047R | | 3230 | 3613 | 7473 | 10228 |
| RET-p.M918T | | 3077 | 3229 | 6877 | 8843 |
| TP53-p.R175H | | 8736 | 10164 | 11559 | 13338 |
| TP53-p.R248Q | | 9132 | 10265 | 12809 | 15163 |
| TP53-p.R273H | | 5733 | 6055 | 8824 | 10315 |
| AKT1-p.E17K | 1 | 5016 | 4629 | 3859 | 3454 |
| APC-p.R1450* | | 5191 | 5621 | 4217 | 4016 |
| ATM-p.C353fs*5 | | 4544 | 4217 | 3732 | 3529 |
| BRAF-p.V600E | | 4929 | 4734 | 3717 | 3390 |
| CTNNB1-p.T41A | | 8879 | 8309 | 6533 | 6554 |
| EGFR-p.D770_N771insG | | 12681 | 12562 | 8960 | 8898 |
| EGFR-p.E746_A750delELREA | | 4603 | 4162 | 3182 | 2964 |
| EGFR-p.L858R | | 6372 | 6641 | 4782 | 4873 |
| EGFR-p.T790M | | 7242 | 7253 | 5343 | 5162 |
| ERBB2-p.A775_G776insYVMA | | 7297 | 7388 | 5237 | 5069 |

| | | | | | |
|---|---|---|---|---|---|
| FLT3-p.D835Y | | 6208 | 6107 | 5090 | 4481 |
| GNA11-p.Q209L | | 6687 | 6158 | 4853 | 4541 |
| GNAQ-p.Q209P | | 5554 | 5034 | 4362 | 3736 |
| GNAS-p.R201C | | 5540 | 5219 | 4384 | 3976 |
| IDH1-p.R132C | | 6889 | 6647 | 5161 | 4727 |
| JAK2-p.V617F | | 5905 | 5639 | 4304 | 4251 |
| KIT-p.D816V | | 6566 | 6242 | 4827 | 4446 |
| KRAS-p.G12D | | 6992 | 6724 | 5501 | 4917 |
| NRAS-p.Q61R | | 5520 | 4798 | 3662 | 3599 |
| PDGFRA-p.D842V | | 6815 | 6309 | 5116 | 4991 |
| PIK3CA-p.H1047R | | 7049 | 6619 | 5623 | 4908 |
| RET-p.M918T | | 5694 | 5324 | 4075 | 3871 |
| TP53-p.R175H | | 7601 | 6987 | 5011 | 4887 |
| TP53-p.R248Q | | 9180 | 8115 | 6079 | 6032 |
| TP53-p.R273H | | 6112 | 5961 | 4465 | 4188 |
